# An intrinsically disordered adaptor protein controls the size and composition of EGFR condensates on membranes

**DOI:** 10.64898/2026.10.15.759085

**Authors:** Maria F. Ullo, Dustin E. Tillman, Hamza Coban, Katherine G. Stratton, Devin D. Baynes, Alfredo Alexander-Katz, Lindsay B. Case

## Abstract

Receptor phase separation is common in signaling pathways, yet mechanisms that limit condensate size and drive their dissolution remain poorly understood. For example, phosphorylated EGFR forms large micron-scale condensates *in vitro*, but assembles into smaller nanoscale clusters in cells, suggesting cellular regulation of condensate size. Here, we show that the intrinsically disordered adaptor protein GAB1 actively dissolves EGFR condensates and re-nucleates smaller, compositionally distinct assemblies in reconstituted supported lipid bilayers and simulations. Dissolution requires unphosphorylated tyrosines within the GAB1 intrinsically disordered region, and tyrosine phosphorylation acts as a regulatory switch that abolishes this activity. GAB1-nucleated assemblies require PH domain-driven oligomerization, which is inhibited by phosphoinositide lipids, linking membrane composition to nucleation. These results uncover two unexpected regulatory features in signaling pathways. First, unphosphorylated tyrosines may function broadly to regulate condensate size or stability, since similar tyrosine-based sequence features are found across hundreds of proteins, including many that function in signal transduction. Additionally, PH domains may play a broader role in protein oligomerization and receptor organization, as an AlphaFold screen reveals dozens of additional PH domains predicted to self-associate. Together, these results establish general mechanisms that function as modular tools for reprogramming signaling through receptor reorganization and may enable new approaches to target such pathways in disease.

## Introduction

Biomolecular condensates formed through phase separation are fundamental organizers of cellular biochemistry. At equilibrium, phase separated condensates grow through coalescence and Ostwald ripening, a phenomenon readily observed *in vitro* with reconstituted condensates growing to tens of microns in size^1^. *In vivo*, condensates are smaller, typically reaching sizes between 100 and 1,000 nm in diamter^2^. The size discrepancy between in vitro condensates and their cellular counterparts can be seen in many systems, including nuclear MED1 transcriptional condensates, cytoplasmic P-bodies, and membrane-associated receptor tyrosine kinase (RTK) condensates^3–5^. This size discrepancy suggests that cells use active processes or distinct molecular mechanisms to restrict condensate size or regulate phase separation. For example, the PD-1 receptor was recently shown to inhibit formation of LAT condensates *in vitro* and in cells^6^. However, the general biophysical mechanisms underlying condensate dissolution or inhibition of condensate growth remain a poorly understood aspect of condensate regulation.

The precise spatiotemporal organization of RTKs, such as the Epidermal Growth Factor Receptor (EGFR), is essential for appropriate activation of downstream signaling. EGFR signaling regulates cell survival, growth, and proliferation; consequently, its mutation or overexpression drives multiple human cancers, including breast cancer^7^. Upon ligand binding, dimerization and autophosphorylation of tyrosine residues in the EGFR C-terminal tail enables the recruitment of multivalent adaptor proteins to propagate downstream RAS/ERK, PI3K/AKT, JAK/STAT, and PLCγ signaling^8^. Effective signaling requires higher-order oligomerization and supramolecular assembly^9^. Several RTKs, including EGFR, can phase separate *in vitro* with downstream adaptors (e.g. SHP2, SHC, GRB2) to form large condensates (1-10 microns)^5^. When anchored on supported lipid bilayers, EGFR phase separates with GRB2 to form membrane-associated micron-sized condensates that increase SOS-dependent Ras activation^10^. However, in cells EGFR forms much smaller, nanometer-sized clusters (∼20 nm)^11^. If phosphorylated EGFR has the intrinsic capacity to form micron-sized condensates, what mechanisms restrict EGFR phase separation in cells? To fully understand EGFR signaling and prevent its oncogenic hyperactivation, we must understand the biophysical basis of EGFR spatial organization.

Because EGFR phase separation depends on multivalent interactions with GRB2^10^, we hypothesized that GRB2-binding proteins might further regulate EGFR spatial organization. GRB2-associated binder-1 (GAB1) is an adaptor protein essential for RTK signaling^12^. GAB1 is comprised of an N-terminal Pleckstrin homology (PH) domain and a 570 amino acid intrinsically disordered region (IDR), and is canonically thought to function as a passive scaffold molecule^13^. In *D. melanogaster* and *C. elegans*, GAB1 orthologs (dos and soc1) are required for RTK signal transduction and function upstream of RAS^14,15^.

In mice, loss of GAB1 is embryonically lethal due to defects in ERK activation^16^. In mammalian EGFR signaling, GAB1 promotes PI3K/AKT signaling and is required for full activation of RAS/ERK^17^. Furthermore, GAB1 is necessary for the transformation of cells with activating mutations of EGFR2, and its overexpression is broadly implicated in the progression, therapeutic resistance, and metastasis of multiple cancers, including breast cancer^18,19^. Despite the central role of GAB1 in both developmental and oncogenic RTK signaling, the fundamental biophysical principles underlying GAB1 function remains an enigma. While the mechanisms of other essential effectors, such as the SOS guanine nucleotide exchange factor, are understood at the atomic level, exactly how the predominantly unstructured GAB1 regulates both PI3K and RAS signaling at the membrane is poorly understood.

In this study, we used *in vitro* reconstitution to investigate how GAB1 regulates EGFR spatiotemporal organization on supported lipid bilayers. We found that GAB1 actively dissolves pre-formed, GRB2-dependent EGFR condensates and nucleates smaller, compositionally distinct EGFR assemblies. In these GAB1-nucleated assemblies, EGFR concentration is drastically reduced, an observation consistent with the spatial anti-correlation of GAB1 and EGFR localization we identify in breast cancer cells. Mechanistically, unphosphorylated tyrosine residues within GAB1’s IDR are required for condensate dissolution, and tyrosine phosphorylation acts as a regulatory switch to abolish this activity. GAB1 has similar tyrosine motifs to the inhibitory adaptor PD-1, which similarly inhibits LAT phase separation. While PD-1 uses these motifs to inhibit LAT receptor signaling, our data suggest that GAB1 uses these motifs to reprogram, rather than inhibit, EGFR signaling. Notably, over 500 additional proteins contain similar motifs in their cytoplasmic IDRs, and these motifs are overrepresented in signaling adaptors and actin binding proteins. Kinetic Monte Carlo simulations that recapitulate EGFR phase separation demonstrate that GAB1 destabilizes EGFR condensates via competitive binding and that GAB1 self-association is required for formation of distinct GAB1 assemblies. Surprisingly, although GAB1 lacks any canonical oligomerization domains, GAB1-nucleated assemblies require the N-terminal PH domain. Using mass photometry, we find that full-length GAB1 forms stable oligomers and the GAB1 PH domain is sufficient for oligomerization. GAB1 oligomerization and the nucleation of GAB1 assemblies are inhibited by phosphoinositide lipid binding, the domain’s canonical function, linking receptor organization to membrane composition. Guided by an AlphaFold screen, we discovered that GAB1 is one of 30 PH domains structurally predicted to dimerize. Thus, a substantial subset of PH domains may function to promote protein oligomerization in addition to their classic lipid-binding roles. While GAB1 has long been recognized as an essential scaffold in RTK signaling networks, our work reveals the distinct biophysical role of this protein as an active remodeler of receptor organization at the membrane and uncovers a generalizable role of cytoplasmic adaptor proteins in controlling the stability, size, and composition of membrane-associated condensates.

## Results

### EGFR phase separates with GRB2 and N-WASP to form micron-sized condensates

To investigate molecular mechanisms regulating the spatiotemporal organization of EGFR, we first aimed to reconstitute EGFR phase separation *in vitro*. Since EGFR is a transmembrane protein and plasma membranes critically facilitate condensate formation^20^, we used supported lipid bilayers (SLBs) as our primary experimental system^21^. To recapitulate intracellular signaling events that occur after EGFR autophosphorylation, we attached the fully phosphorylated, His-SUMO-tagged cytoplasmic tail of EGFR (“EGFR”) to POPC SLBs containing 2% Ni-NTA lipids (Fig. 1A). As in previous studies^10^, the SUMO domain functions as a spacer between the surface of the SLB and the tail of EGFR, where the native kinase domain would normally be positioned. Previous work identified that the phosphorylated EGFR cytoplasmic tail, which is highly disordered, only forms condensates in vitro when an SH2-containing adaptor protein (e.g. GRB2, SHP2, SHC) is present^5,10^. When we incubated Alexa488-labeled EGFR with GRB2 and the BPVCA domain of N-WASP (“N-WASP”), we observed rapid formation of micron-sized condensates, similar to what has previously been reported (Fig. 1B).

**Figure 1:**
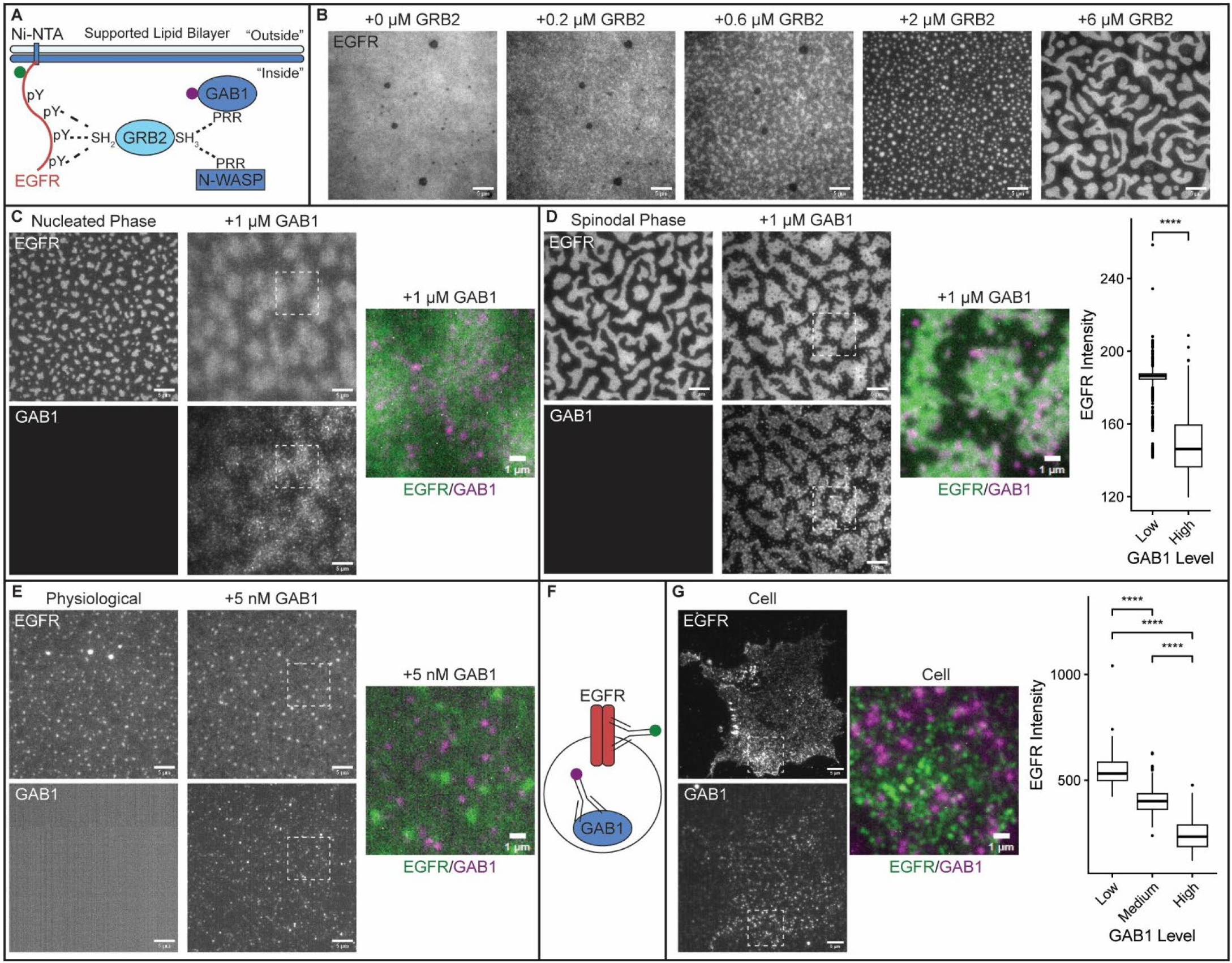
GAB1 destabilizes EGFR condensates and nucleates distinct assemblies. **(A)** His-tagged, phosphorylated EGFR cytoplasmic tails with Alexa488 and a SUMO spacer (“EGFR”) are attached to POPC SLBs with 2% Ni-NTA lipids and incubated with distinct EGFR adaptor proteins. Dashed lines indicate established interactions. **(B)** EGFR undergoes GRB2-dependent phase separation on SLBs when the BPVCA domain of N-WASP (“N-WASP”) is present. **(C)** GAB1 dissolves pre-formed EGFR condensates formed through nucleation on SLBs. LEFT: Images before GAB1 is added. RIGHT: Images one minute after adding GAB1. **(D)** GAB1 destabilizes EGFR condensates formed through spinodal decomposition on SLBs. LEFT: Images before adding GAB1. RIGHT: Images one minute after adding GAB1. Box plot shows quantification of EGFR intensity in clusters grouped by low or high GAB1 intensity. **(E)** EGFR localization on SLBs using physiological concentrations of all proteins. LEFT: Images with EGFR/GRB2/N-WASP. RIGHT: Images one minute after adding GAB1. **(F)** Endogenous EGFR and GAB1 are visualized in fixed MDA-MB-468 cells via immunofluorescence. **(G)** EGFR and GAB1 form distinct clusters in cells. Box plot shows quantification of EGFR intensity in clusters grouped by low, medium, or high GAB1 intensity. Concentrations: 150 nM EGFR (B-D), 1 μM N-WASP (B-D), 2 μM GRB2 (C), 6 μM GRB2 (D), 1 μM GAB1 (C, D); Physiological concentrations (E): 114 nM EGFR, 662 nM GRB2, 138 nM N-WASP, 5 nM GAB1. SLBs: 97.9% POPC, 2% Ni-NTA, 0.1% DOPE-PEG5000. In C-G the enlarged pseudocolored image corresponds to the region denoted by the white dashed box. Scale bars: 5 μm; Inset scale bars: 1 μm. Statistics: two-sided Student’s t-test (****: p < 0.0001; n > 188 condensates per condition).

EGFR condensates are GRB2-dependent, and we observe two distinct regimes of EGFR phase separation. With lower GRB2 concentration (≤ 2 μM) EGFR phase separates through nucleation into discrete puncta (Fig. 1B). With higher GRB2 concentrations (≥ 6 μM), EGFR phase separates through spinodal decomposition (Fig. 1B), a well-characterized phenomenon that occurs when there is no thermodynamic barrier for phase separation^22^. In addition to GRB2, condensate formation required a third adaptor protein containing protein rich regions (PRR), such as N-WASP (Fig. 1B) or SOS (Supp. Fig. 1A). Thus, EGFR phase separation is driven by multivalent pY-SH2-SH3-PRR interaction networks, similar to what has been observed with LAT and nephrin receptors^23,24^. Since N-WASP formed larger EGFR condensates than SOS at similar concentrations, we used N-WASP in all experiments. To orthogonally validate EGFR phase separation with GRB2 and N-WASP, we also reconstituted EGFR condensates as droplets in solution and on giant unilamellar vesicle (GUV) membranes (Supp. Fig. 1B-1E). Together, these results demonstrate that multivalent protein interactions between EGFR, GRB2, and N-WASP promote phase separation across experimental systems.

### GAB1 destabilizes EGFR condensates and nucleates distinct EGFR assemblies

To identify molecules that potentially control GRB2-mediated EGFR phase separation, we analyzed features of proteins that interact with EGFR and GRB2. We hypothesized that ideal regulators would possess long cytoplasmic IDRs, providing tunable multi-valency, and many phosphorylation-accessible tyrosines, enabling toggled control by kinases and phosphatases. To characterize the 28 proteins that directly interact with both EGFR and GRB2, as defined by the Human Protein Atlas, we used DeepTMHMM to predict cytoplasmic domains, AIUPred to predict disordered regions, and Musite to predict phosphorylation-accessible tyrosines^25–28^. Top candidates included EGFR paralogs (EGFR2/3/4), which form heteromeric dimers with EGFR^29^, as well as LAT and FAK, which form condensates in other systems^23,30–34^ (Supp. Fig. 1F). In contrast, SOS, GRB2, N-WASP, and PI3K have short cytoplasmic IDRs with few phosphorylation-accessible tyrosines. Notably, GAB1, an essential adaptor protein that activates PI3K/AKT signaling downstream of EGFR^35,36^, was the highest-scoring candidate. Although GAB1 is co-enriched with GRB2 in some fusion oncoprotein condensates^37,38^, nothing is known about how GAB1 might control formation, stability, and/or biophysical properties of EGFR condensates.

To investigate how GAB1 impacts GRB2-dependent EGFR phase separation, we added Alexa647-labeled GAB1 to pre-formed EGFR/GRB2/N-WASP condensates on SLBs. Strikingly, GAB1 rapidly dissolves condensates formed through nucleation and destabilizes the condensed phase of condensates formed through spinodal decomposition (Fig. 1C, 1D). Furthermore, after adding GAB1, small GAB1 clusters nucleate *de novo* (Fig. 1C, 1D). These clusters, which we refer to as “GAB1 assemblies” to differentiate them from the initial EGFR condensates, are enriched in GAB1 and have lower levels of EGFR compared to the initial EGFR condensates. This results in a significant anti-correlation between EGFR and GAB1 intensities, with the highest EGFR intensities occurring where GAB1 is not enriched (Fig. 1D). We observe similar effects of GAB1 on EGFR condensates in droplet assays and on GUV membranes (Supp. Fig. 1G, 1H), demonstrating that GAB1’s ability to destabilize EGFR condensates and nucleate distinct GAB1 assemblies does not require membranes and is not an artifact of SLBs. Finally, when we combine physiological concentrations of EGFR, GRB2, N-WASP, and GAB1 on SLBs^39^, we observe the co-existence of small EGFR-rich clusters with small GAB1-rich clusters (Fig. 1E). From these experiments, we conclude that GAB1 is sufficient to regulate GRB2-dependent EGFR phase separation and can induce dramatic changes in the spatial organization of EGFR on membranes.

We next examined the spatial organization of GAB1 and EGFR in cells. We performed immunostaining of endogenous EGFR and GAB1 in MDA-MB-468 cells, a human triple-negative breast cancer cell line with EGFR amplification (Fig. 1F). MDA-MB-468 cells contain high levels of EGFR1, no EGFR2, and low, baseline levels of EGFR3 and EGFR4^40^. In cells, we observe the co-existence of small EGFR-rich clusters with small GAB1-rich clusters (Fig. 1F), strikingly similar to what we observe with physiological protein concentrations *in vitro* (Fig. 1E). Clusters with low GAB1 intensity have significantly higher EGFR intensities than clusters with medium or high GAB1 intensity (Fig. 1G). Thus, GAB1 intensity is anti-correlated with EGFR intensity both *in vitro* and in cells. In cells, this anti-correlation is weaker after serum starvation, which inhibits EGFR activity and clustering^41,42^, and is quickly restored after addition of EGF (Supp. Fig. 1I). Together, these data demonstrate that our *in vitro* reconstitution on SLBs is sufficient to recapitulate the organization of GAB1 and EGFR on cellular plasma membranes and suggest that this organization dynamically changes downstream of receptor activation. Thus, we next used the *in vitro* SLB assay to dissect molecular features underlying GAB1’s ability to reshape EGFR organization.

### The PH domain and IDR of GAB1 mediate distinct functions of GAB1

To elucidate how GAB1 destabilizes EGFR condensates and nucleates distinct GAB1 assemblies, we first investigated functions of different GAB1 domains (Fig. 2A). GAB1 has a N-terminal PH domain thought to bind PIP3 lipids to facilitate GAB1-dependent PI3K signaling^13,43,44^. While the PH domain is the only structured domain, the 578 residue C-terminal IDR contains a PRR with two GRB2 binding sites^45,46^. To test functions of these GAB1 domains on EGFR and GAB1 organization, we incubated pre-formed EGFR/GRB2/N-WASP condensates on SLBs with either GAB1-WT (residues 1-694), GAB1 lacking its PH domain (GAB1-ΔPH; 116-694), the GAB1 PRR (GAB1-PRR; 331-530), or a segment of the GAB1 IDR (GAB1-IDR493; 493-694). As expected, GAB1-WT destabilizes EGFR condensates and nucleates distinct GAB1 assemblies (Fig. 2B; Supp. Fig. 2A). GAB1-ΔPH also destabilizes EGFR condensates, but nucleates far fewer GAB1 assemblies. Although GAB1-IDR493, which contains one GRB2 binding site, weakly destabilizes EGFR condensates, it does not nucleate GAB1 assemblies, and GAB1-PRR neither destabilizes EGFR condensates nor nucleates new GAB1 assemblies. Additionally, the PRR of SOS (SOS-PRR; residues 1117-1319; 2 GRB2 binding sites^47^) does not disrupt EGFR condensates (Supp. Fig. 2B). Thus, GRB2 binding sites are not sufficient for destabilization of EGFR condensates. Together, these findings demonstrate that the PH domain contributes to the nucleation of GAB1 assemblies while the entire GAB1 IDR is required to destabilize EGFR condensates.

**Figure 2:**
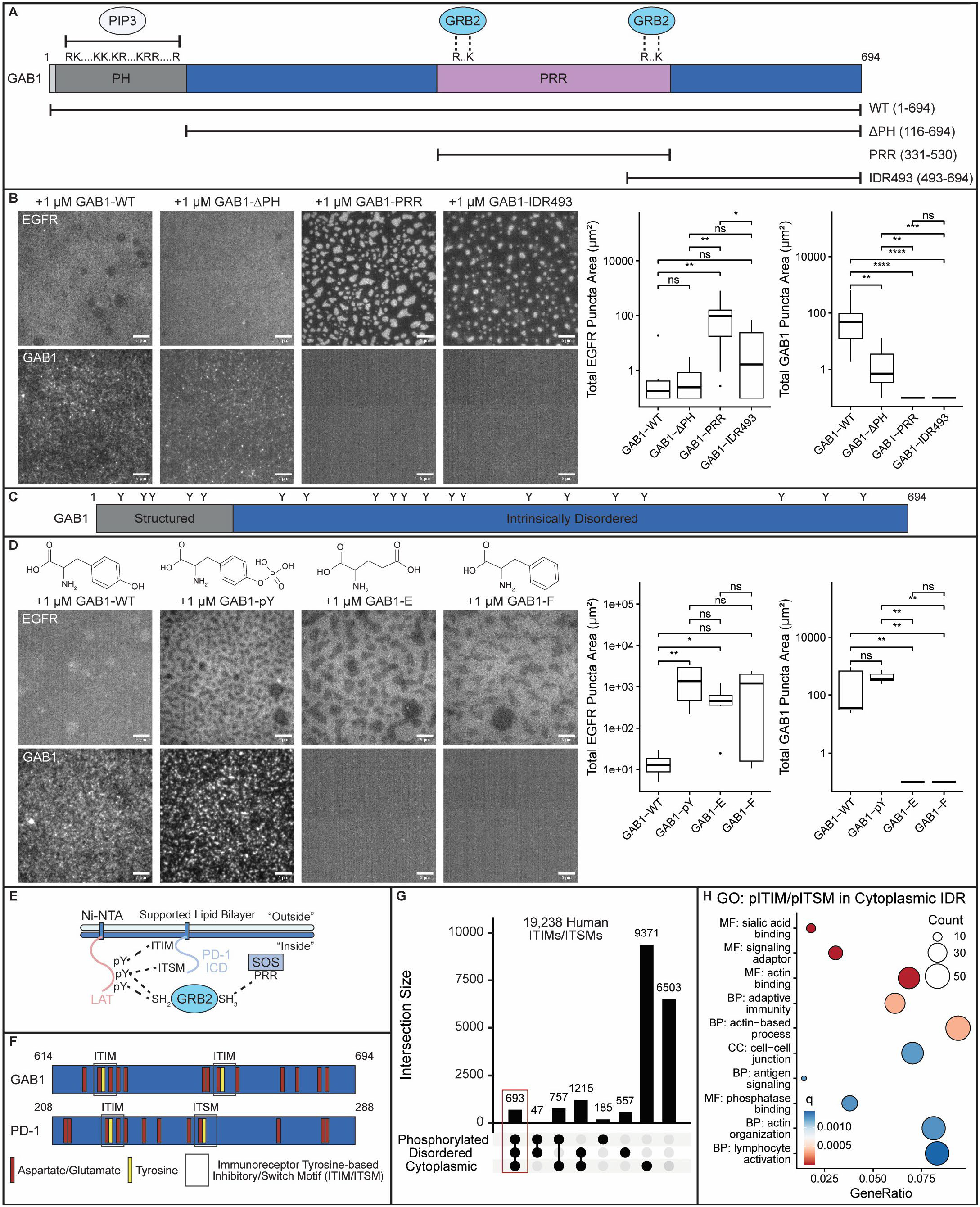
The PH domain and IDR of GAB1 regulate distinct aspects of EGFR and GAB1 spatial organization. **(A)** GAB1 has a structured PH domain that binds PIP3 followed by a long IDR, which contains a PRR with two GRB2 binding sites. **(B)** Images of EGFR condensates and GAB1 assemblies on SLBs after adding the following GAB1 variants: Full-length GAB1 (GAB1-WT), GAB1 without its PH domain (GAB1-ΔPH), the GAB1 PRR (GAB1-PRR), a C-terminal segment of the IDR (GAB1-IDR493). Box plots show quantification of total EGFR puncta area and total GAB1 puncta area. **(C)** GAB1 has twenty tyrosines across its structured and disordered regions – all nine phosphorylation-accessible tyrosines are in the IDR. **(D)** Images of EGFR condensates and GAB1 assemblies on SLBs after adding the following full-length GAB1 variants: unphosphorylated GAB1 (GAB1-WT), fully phosphorylated GAB1 (GAB1-pY), GAB1 with all 20 tyrosines mutated to either glutamic acid (GAB1-E) or phenylalanine (GAB1-F). Box plots show quantification of total EGFR puncta area and total GAB1 puncta area. **(E)** Recent work identified that incubating pLAT with the intracellular domain (ICD) of PD-1 inhibits formation of LAT/GRB2/SOS condensates on SLBs via the unphosphorylated tyrosines in its immunoreceptor tyrosine-based inhibitory/switch motifs (ITIMs/ITSMs)^6^. Dashed lines indicate established interactions. **(F)** The last 80 residues of GAB1 and PD-1 each have two ITIMs/ITSMs with similar spacing in acidic patches. These features underlie inhibition of LAT condensate formation by PD-1. **(G)** Upset plot of all human ITIMs/ITSMs based on their phosphorylation accessibility and location in a cytoplasmic IDR. The 693 ITIMs/ITSMs in 588 proteins that were selected for GO analysis are indicated with a red box. **(H)** The top ten enriched GO terms for proteins with at least one phosphorylation-accessible ITIM/ITSM in a cytoplasmic IDR relative to all proteins with at least one ITIM/ITSM. Terms are colored based on their q-value, sized based on their number of proteins, and preceded by their GO “aspects” (MF: molecular function; BP: biological process; CC: cellular component). Concentrations for all experiments: 150 nM EGFR, 1 μM N-WASP, 6 μM GRB2, 1 μM GAB1 variant; SLBs: 97.9% POPC, 2% Ni-NTA, 0.1% DOPE-PEG5000. Scale bars: 5 μm. Statistics: two-sided Wilcoxon test (****: p < 0.0001; ***: p < 0.001; **: p < 0.01; *: p < 0.05; ns: p > 0.05; n > 3 replicates

To determine features within the GAB1 IDR that might contribute to GAB1 nucleation or EGFR condensate destabilization, we next investigated GAB1’s 20 tyrosines (Fig. 2C). Tyrosine residues are unique stickers within IDRs, due to their ability to engage in pi-pi stacking and pi-cation interactions through the aromatic ring as well as hydrogen bonding through the hydroxyl group^48^. Thus, tyrosines distributed along IDRs often mediate multivalent IDR-IDR interactions^49^. Specific phosphatases and RTKs (including EGFR) control the phosphorylation state of GAB1’s tyrosines to regulate adaptor protein binding^50,51^, and the nine phosphorylation-accessible tyrosines are all in GAB1’s cytoplasmic IDR (Supp. Fig. 1F).

To determine the importance of tyrosines in GAB1’s ability to reshape EGFR organization, we incubated preformed EGFR/GRB2/N-WASP condensates on SLBs with either wild-type, unphosphorylated GAB1 (GAB1-WT), fully tyrosine-phosphorylated GAB1 (GAB1-pY), or GAB1 with all tyrosines mutated to either glutamic acid (GAB1-E; phospho-mimic) or phenylalanine (GAB1-F; aromatic mimic). As expected, GAB1-WT destabilizes EGFR condensates and nucleates distinct GAB1 assemblies (Fig. 2D; Supp. Fig. 2A). Intriguingly, GAB1-pY nucleates GAB1 assemblies but fails to destabilize EGFR condensates, suggesting that tyrosine phosphorylation regulates GAB1-dependent changes to EGFR organization. In contrast, neither GAB1-E nor GAB1-F destabilize EGFR condensates or nucleate GAB1 assemblies, indicating that tyrosine-specific side chain interactions are essential for both functions of GAB1. These results identify GAB1 tyrosines as important mediators of GAB1 assembly nucleation, as well as toggleable regulators of EGFR condensate stability.

Similar to EGFR, phosphorylated LAT forms condensates with GRB2 and SOS^23^. LAT condensates promote T cell activation, and recent work identified that the intracellular domain of the immune inhibitory receptor PD-1 inhibits the formation of LAT condensates via unphosphorylated tyrosines (Fig. 2E)^6^. Similar to GAB1, mutation of the PD-1 tyrosines to phenylalanine abolishes the ability of the protein to inhibit receptor phase separation^6^. Although no distinct PD-1 assemblies are described, these results are strikingly similar to destabilization of EGFR condensates by GAB1’s unphosphorylated tyrosines (Fig. 2D), suggesting a potentially generalizable mechanism for inhibition of phase separation. Indeed, the final 80 residues of GAB1 and PD-1 both contain two immunoreceptor tyrosine-based inhibitory/switch motifs (ITIMs/ITSMs) in acidic patches, the critical feature contributing to PD-1’s inhibition of LAT condensate formation (Fig. 2F; Supp. Fig. 2C, 2D)^6^. Since GAB1-IDR493 does not fully dissolve EGFR condensates, the GAB1 ITIMs likely function with the PRR to destabilize EGFR condensates.

To identify other proteins with these condensate-destabilizing sequence features, we analyzed all 19,238 ITIMs/ITSMs in the human proteome and selected the 693 that are phosphorylation-accessible and in cytoplasmic IDRs (Fig. 2G), resulting in 588 candidate proteins. Among these candidates, gene ontology (GO) analysis identifies overrepresentation of signaling adaptors (e.g. GAB1-4, IRS1-2, FRS2-3, SHANK2-3, MAPK8IP1-2, PAG1, DDX3X, DOK2, NLRP1,) actin binders (e.g. ABLIM1-3, MYO5A-7B, CEACAM1, SHANK3, WAS, MAP1B, TLN1, TNS1, VASP, VCL), adaptive immunity proteins (e.g. PD-1, RAP1GAP1-2, LLIRB1-5, SLAMF6-7, CEACAM1, PAG1, WAS, AIRE, SIT1, NFKB2), and other GO terms (Fig. 2H). Thus, these diverse proteins might inhibit formation of specific condensates with functions beyond immunity, including in cytoskeletal remodeling, cell adhesion, and signal transduction.

### In silico reconstitution of EGFR phase separation predicts that GAB1 oligomers form stable assemblies

To investigate biophysical principles driving EGFR phase separation, GAB1-mediated condensate destabilization, and nucleation of *de novo* GAB1 assemblies, we reconstituted our system *in silico*. Briefly, we simulated the membrane as a minimal, two-dimensional system in equilibrium with a fluid reservoir above the bilayer containing free proteins. To model this system, we employed Kinetic Monte Carlo (KMC), enabling explicit incorporation of binding reactions and protein states. We assumed that EGFR/GRB2 complexes are permanent species and simulated stochastic exchange with the reservoir containing N-WASP and GAB1 (Supp. Fig. 3A-3C). We included oligomerized GAB1 as a distinct species that exchanges between being bound on the membrane or free in solution using a phenomenological self-limiting droplet free energy^52,53^.

**Figure 3:**
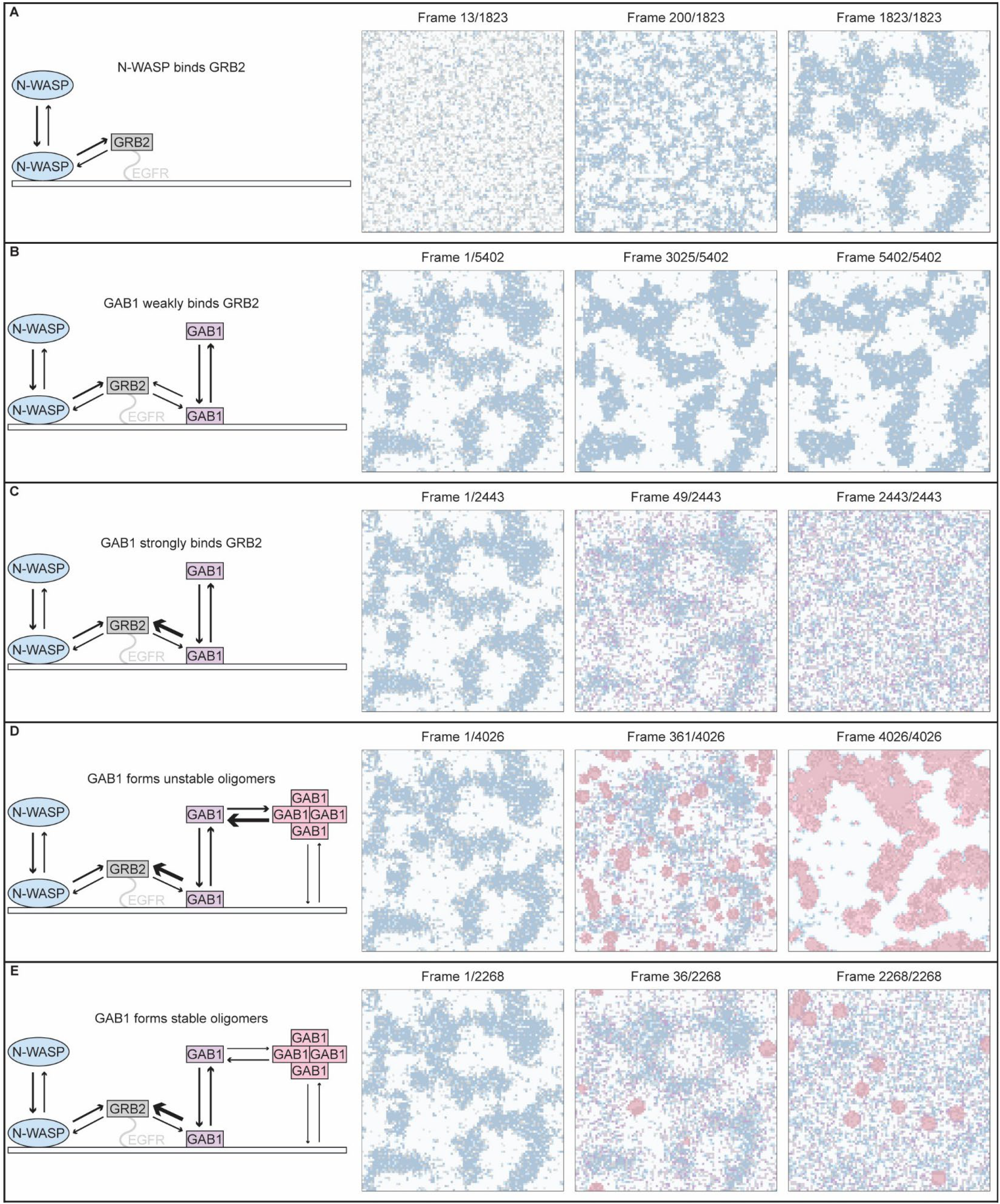
*In silico* reconstitution of EGFR phase separation predicts that GAB1 oligomers form stable assemblies. **(A)** Simulation of EGFR condensate formation. N-WASP in solution exchanges with membrane-diffusing N-WASP, which binds EGFR/GRB2 complexes, resulting in spinodal decomposition of EGFR. **(B-E)** Simulation of GAB1 added to EGFR condensates with different GAB1 parameters. **(B)** GAB1 in solution exchanges with membrane-diffusing GAB1, which weakly binds EGFR/GRB2 complexes, leading to no effect on EGFR condensates. **(C)** Strengthening the interaction between GAB1 and EGFR/GRB2 leads to destabilization of EGFR condensates, but no GAB1 nucleation. **(D)** Labile GAB1 oligomers with rapid monomer/oligomer exchange destabilize EGFR condensates and form a large spinodal phase. **(E)** Stable GAB1 oligomers with slow monomer/oligomer exchange destabilize EGFR condensates and nucleate distinct GAB1 puncta. All parameters are in Supplemental Figure 3.

We first aimed to reconstitute EGFR/GRB2/N-WASP phase separation *in silico*. We omitted GAB1 from these simulations while modeling bulk/membrane exchange of N-WASP and direct, multivalent binding of membrane-diffusing N-WASP with EGFR/GRB2 complexes (Fig. 3A). As with our *in vitro* reconstitution (Fig. 1B), we identify spinodal decomposition of EGFR/GRB2/N-WASP under these conditions, demonstrating how EGFR phase separation is facilitated by the multivalent N-WASP PRR. We next simulated the addition of GAB1 to pre-formed EGFR condensates by modeling bulk/membrane exchange of GAB1 and direct, monovalent binding of membrane-diffusing GAB1 with EGFR/GRB2 complexes (Fig. 3B). When we model GAB1 as a weaker binder of EGFR/GRB2 than N-WASP, there is no change to EGFR condensates. However, when we model GAB1 as a stronger binder of EGFR/GRB2 than N-WASP, EGFR condensates are rapidly destabilized, but GAB1 does not form puncta (Fig. 3C), an outcome similar to our *in vitro* addition of GAB1-ΔPH to pre-formed EGFR condensates (Fig. 2B). These results identify that competitive displacement of N-WASP by GAB1 via its tyrosine-rich IDR, similar to PD-1’s inhibition of LAT condensate formation via polyphasic linkage^6^, is sufficient for destabilization of EGFR condensates, but is not sufficient to nucleate GAB1 assemblies.

To investigate how GAB1 forms *de novo* assemblies, we next considered the possibility that GAB1 oligomerizes in solution. Oligomerization increases molecular valency and homooligomerization often regulates protein phase separation^54–57^. Exchange kinetics are a key oligomerization parameter – fast exchange kinetics lead to “labile” oligomers, while slow exchange kinetics result in “stable” oligomers. We first modeled labile GAB1 oligomers with fast formation and dissociation kinetics (Fig. 3D). Under these conditions, GAB1 destabilizes EGFR condensates, but rapidly promotes the phase separation of a large EGFR-rich phase, effectively replacing N-WASP in the EGFR spinodal regime. We next modeled stable GAB1 oligomers with slow formation and dissociation kinetics (Fig. 3E). Under these conditions, GAB1 destabilizes EGFR condensates and nucleates distinct GAB1 puncta, recapitulating our *in vitro* experiments with GAB1-WT (Fig. 1C-1E). Together, these *in silico* simulations reconstitute GRB2-dependent EGFR phase separation, demonstrate that GAB1 competition with N-WASP destabilizes EGFR condensates, and predict that the nucleation of GAB1 assemblies requires the formation of stable, slowly exchanging GAB1 homooligomers.

### The GAB1 PH domain mediates formation of stable GAB1 oligomers

To test our *in silico* prediction that GAB1 forms stable assemblies with slow formation and dissociation kinetics, we performed fluorescence recovery after photobleaching (FRAP) experiments on SLB condensates with distinct compositions. We first induced phase separation of EGFR, Alexa568-labeled GRB2, and N-WASP on SLBs, bleached GRB2 in nucleated condensates, and quantified fluorescence recovery (Fig. 4A). We then added Alexa647-labeled GAB1, bleached in either the GRB2 or GAB1 channel, and quantified fluorescence recovery (Fig. 4B, 4C). While GRB2 is extremely mobile in both conditions (∼100% mobile fraction), GAB1 is highly immobile (∼8% mobile fraction), demonstrating that GAB1 assemblies are quite stable with few GAB1 molecules dissociating over the 30 second FRAP time course (Fig. 4D, 4E). GRB2 and GAB1 have similar t_1/2_ values, and GRB2 t_1/2_ is higher in the absence of GAB1, suggesting that these proteins might co-diffuse as a stable complex (Supp. Fig. 4A). These results confirm that GAB1 assemblies are more stable than EGFR condensates, consistent with the distinct, persistent GAB1-rich state identified in our KMC simulations.

**Figure 4:**
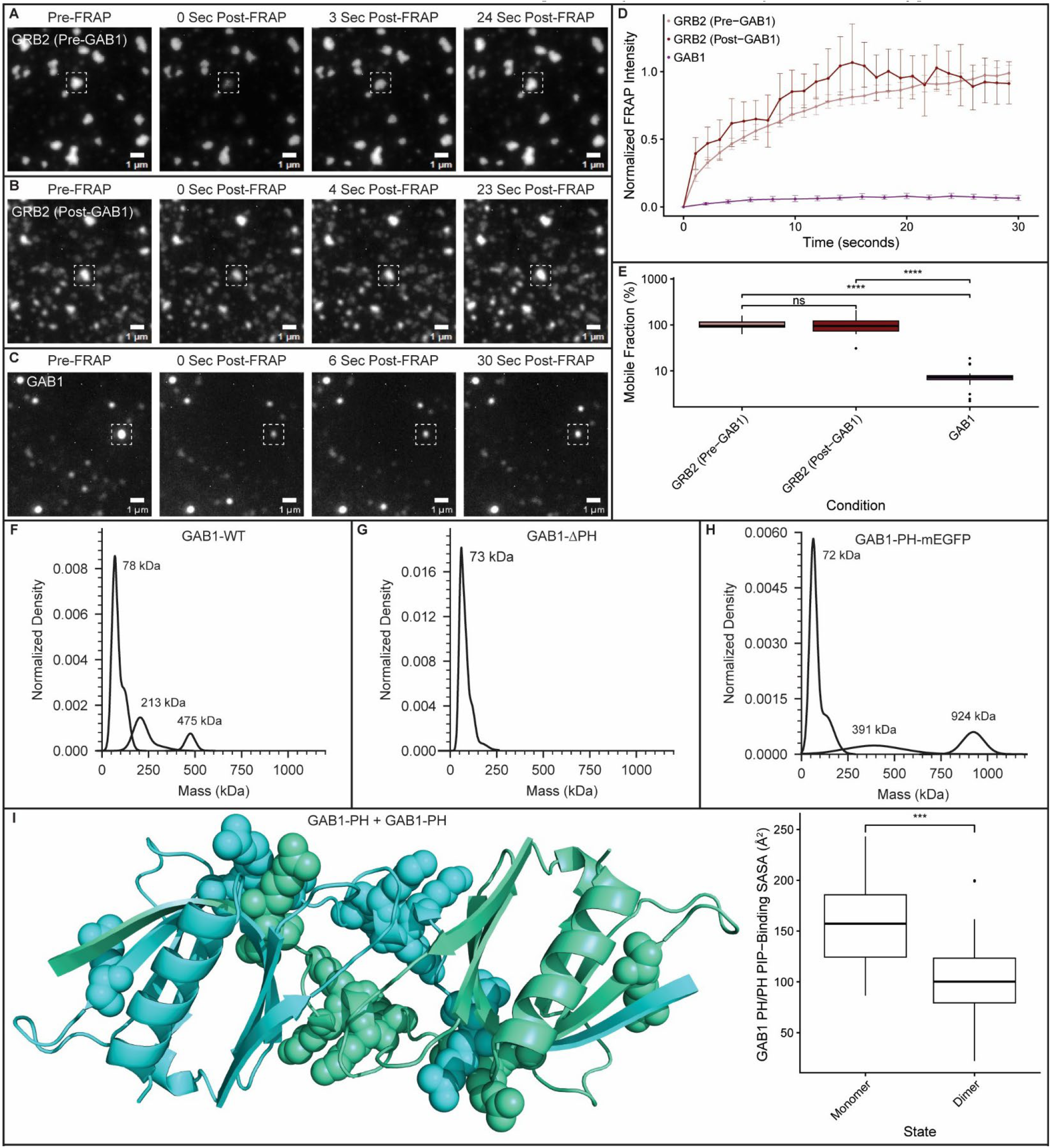
The GAB1 PH domain promotes the formation of stable GAB1 oligomers. **(A)** Fluorescence Recovery After Photobleaching (FRAP) analysis of GRB2 within EGFR/GRB2/N-WASP puncta enables quantification of condensate properties on SLBs. **(B)** Adding GAB1 destabilizes EGFR condensates and enables FRAP analysis of GRB2 within GAB1/GRB2 puncta. **(C)** FRAP analysis of GAB1 within GAB1/GRB2 assemblies enables quantification of GAB1 mobility. **(D)** Quantification of FRAP analysis from A-C. Error bars are 95% confidence intervals (n > 16 condensates per condition). **(E)** Mobile fraction calculated from A-C. **(F-H)** Mass photometry measurements to determine the mass of purified GAB1 variants in solution at 20 nM. **(F)** Wild-type GAB1 (GAB1-WT). **(G)** GAB1 without its PH domain (GAB1-ΔPH). **(H)** The PH domain of GAB1 fused to mEGFP (GAB1-PH-mEGFP). **(I)** *In silico* modeling with AlphaFold predicts a GAB1-PH dimer. GAB1 PIP-binding residues are spheres. The box plot displays the solvent-accessible surface area (SASA) of these PIP-binding residues in the predicted monomer and dimer structures. Concentrations: 150 nM EGFR (A-C), 1 μM N-WASP (A-C), 6 μM GRB2 (A-C), 1 μM GAB1 (A-C), 20 nM GAB1 variants (F-H); SLBs: 97.9% POPC, 2% Ni-NTA, 0.1% DOPE-PEG5000. Statistics: two-sided Wilcoxon test (****: p < 0.0001; ***: p < 0.001; ns: p > 0.05; n = 10 PIP-binding residues per chain). Scale bars: 1 μm.

Our simulations predict that persistent GAB1 self-association could underly the formation of stable GAB1 assemblies. To investigate if GAB1 forms stable oligomers in solution, we performed mass photometry to determine the molecular weight of species in a solution of 20 nM GAB1-WT. We identify a monomeric GAB1-WT mass photometry peak (∼78 kDa), as well as peaks that correspond to GAB1-WT oligomers (∼213 kDa, 475 kDa), suggesting that GAB1 oligomerizes with a K_d_ < 200 nM^58^ (Fig. 4F). We confirmed the ability of GAB1-WT to form higher-order oligomers by combining 1 μM Alexa647-labeled GAB1-WT in a buffer with crowding agent (5% PEG) and imaging with fluorescence microscopy. GAB1-WT forms micron-sized, oligomeric assemblies in solution (Supp. Fig. 4B). However, these PEG-induced complexes are distinct from the nucleated GAB1-WT assemblies that we observe on SLBs in the presence of EGFR, GRB2, and N-WASP (Fig. 1C-1E). These findings validate our *in silico* prediction that GAB1 forms stable oligomers in solution.

Our simulations predict that stable GAB1 oligomers enable the nucleation of GAB1 assemblies on SLBs.

Although GAB1 lacks a canonical oligomerization domain, the PH domain is required for robust nucleation of GAB1 assemblies (Fig 2D). Furthermore, PH domains have been reported to promote the homodimerization of two other proteins (PDPK1 and BTK)^59–62^. Thus, we hypothesized that PH domain self-association could underly GAB1 oligomerization. To test this, we performed mass photometry of GAB1-ΔPH and the isolated GAB1 PH domain fused to monomeric EGFP (GAB1-PH-mEGFP). Mass photometry can only detect masses over 30 kDa, so the mEGFP fusion is required to analyze isolated PH domains. GAB1-ΔPH has a single, monomeric mass photometry peak (∼73 kDa; Fig. 4G), demonstrating that the PH domain is necessary for GAB1 oligomerization. In contrast, GAB1-PH-mEGFP has a monomeric peak (∼72 KDa) as well as several peaks corresponding to higher-order oligomers (∼397 kDa, 924 kDa; Fig. 4H). These data demonstrate that the GAB1 PH domain is necessary and sufficient for oligomerization.

We orthogonally verified these results by combining 1 μM Alexa647-labeled GAB1-ΔPH or 1 μM GAB1-PH-mEGFP in a buffer containing crowding agent (5% PEG). GAB1-PH-mEGFP formed micron-sized, oligomeric complexes in solution, confirming the PH domain is sufficient for oligomerization (Supp. Fig. 4B). GAB1-ΔPH formed much smaller assemblies than GAB1-WT, suggesting that the IDR contains some capacity to self-associate at high concentrations (Supp. Fig. 4B). This result is consistent with our observation that GAB1-ΔPH dramatically reduces, but does not eliminate, GAB1-nucleated assemblies on SLBs (Fig. 2B). Neither GAB1-PRR nor GAB1-IDR493, both lacking the GAB1 PH domain, form PEG-induced assemblies in solution (Supp. Fig. 4B), consistent with their inability to nucleate GAB1 assemblies on SLBs (Fig. 2B). Additionally, GAB1-pY, GAB1-E, and GAB1-F fail to form PEG-induced assemblies in solution (Supp. Fig. 4B), suggesting that tyrosine-specific side chain interactions within the IDR contribute to GAB1 oligomerization. Together, these results identify the GAB1 PH domain as the major driver of GAB1 oligomerization and suggest that GAB1 can make additional IDR-mediated intermolecular interactions that are too weak to detect with mass photometry.

To investigate functional consequences of GAB1 oligomerization, we computationally predicted structures of GAB1-PH homodimers with AlphaFold^63^. Notably, no structures of GAB1 have been solved, potentially impacting the quality of AlphaFold-based predictions^64^. Nevertheless, we identify a GAB1-PH homodimer structure with low to moderate confidence (Fig. 4I; Supp. Fig. 4C), as quantified by several common scoring metrics^63,65,66^ (ipTM = 0.34; pDockQ = 0.29; AlphaJudge Interface Score = 142), in line with our experimental observation that GAB1-PH self-associates (Fig. 4H). Since the canonical function of PH domains is to bind membranes containing phosphoinositide lipids^67^, and since the GAB1 PH domain binds PIP3^13^, we investigated if the GAB1-PH PIP-binding residues are solvent-accessible in the modeled homodimer state^68^. Strikingly, the solvent-accessible surface area (SASA) of GAB1-PH PIP-binding residues is predicted to significantly decrease in GAB1-PH homodimers (Fig. 4I). These *in silico* findings support our experimental identification of PH-mediated GAB1 oligomerization and suggest that mammalian GAB1 monomers might bind PIPs more strongly than GAB1 oligomers.

### PIPs negatively regulate GAB1 oligomerization and function

To experimentally test if PIPs influence the spatial organization of EGFR and GAB1, we employed GUVs, which minimize artifacts caused by interactions between charged PIP headgroups and coverslip glass^69^. We generated POPC GUVs with Ni-NTA lipids and either 0% PIPs (“POPC GUVs”), 2% PIP2 (“PIP2 GUVs”), or 2% PIP3 (“PIP3 GUVs”), then added Alexa488-labeled EGFR. We quantified the squared coefficient of variation (CV^2^) of EGFR fluorescence along GUV surfaces, which increases with EGFR clustering, to validate that EGFR binding to GUVs is not impacted by PIPs (Fig. 5A; Supp. Fig. 5A-5D). We incubated these GUVs with GRB2 and N-WASP and observe condensate formation on GUV surfaces across conditions (Supp. Fig. 5B), as we previously observed on POPC GUVs (Supp. Fig. 1E). These results demonstrate that GRB2-dependent EGFR phase separation on GUVs does not depend on the membrane PIP composition.

**Figure 5:**
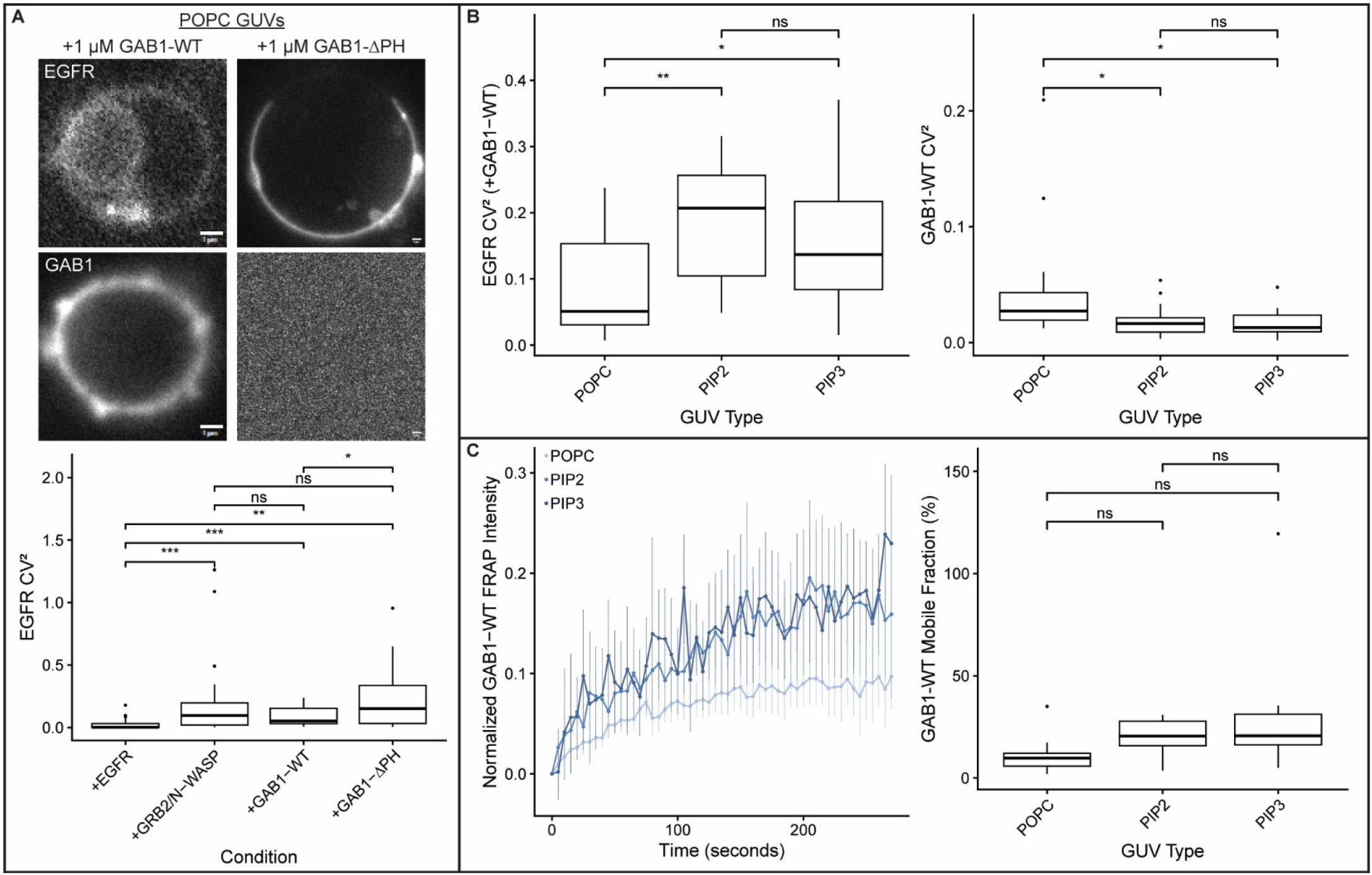
Phosphatidylinositol phosphate lipids negatively regulate GAB1 oligomerization and function. **(A)** Images showing wild-type GAB1 (GAB1-WT) or GAB1 lacking its PH domain (GAB1-ΔPH) added to pre-formed EGFR condensates on GUVs containing no phosphatidylinositol phosphate lipids (“POPC GUVs”). GAB1-WT, but not GAB1-ΔPH, is recruited to GUV surfaces, nucleates distinct assemblies, and destabilizes EGFR condensates. Box plots quantify the squared coefficient of variation (CV2) of EGFR fluorescence along GUV surfaces, which increases with clustering. **(B)** Quantification of CV^2^ of EGFR and GAB1 fluorescence along GUV surfaces after adding GAB1-WT to pre-formed EGFR condensates on POPC GUVs, GUVs containing 2% phosphatidylinositol 4,5-bisphosphate (“PIP2 GUVs”), or GUVs containing 2% phosphatidylinositol (3,4,5)-trisphosphate (“PIP3 GUVs”). **(C)** Fluorescence recovery after photobleaching (FRAP) analysis of GAB1 assemblies on GUV surfaces with distinct compositions. Concentrations: 150 nM EGFR, 1 μM N-WASP, 6 μM GRB2, 1 μM GAB1 construct; POPC GUVs: 97.9% POPC, 2% Ni-NTA, 0.1% DOPE-PEG5000; PIP2 GUVs: 95.9% POPC, 2% Ni-NTA, 2% PIP2, 0.1% DOPE-PEG5000; PIP3 GUVs: 95.9% POPC, 2% Ni-NTA, 2% PIP3, 0.1% DOPE-PEG5000.

To investigate if PIPs impact GAB1’s ability to reshape EGFR organization, we added Alexa647-labeled GAB1-WT or GAB1-ΔPH to PIP-containing GUVs with pre-formed EGFR condensates. GAB1-WT forms more puncta on POPC GUVs relative to PIP2 or PIP3 GUVs (Fig. 5B). Since puncta formation requires stable GAB1 oligomers, these data suggest that the presence of PIPs favors recruitment of PIP-binding, monomeric GAB1 over the oligomeric GAB1 with its PIP-binding residues buried (Fig. 4I). Across all GUV types, GAB1-WT mildly destabilizes EGFR condensates (Fig. 5A; Supp. Fig. 5C, 5D) but this effect is strongest on POPC GUVs (Fig. 5B), suggesting that GAB1 oligomers more effectively destabilize EGFR condensates. GAB1-ΔPH is not recruited to the surface of GUVs (Fig. 5A; Supp. Fig. 5C, 5D), regardless of PIP content. Thus, the PH domain is required to efficiently recruit GAB1 to the membrane surface independent of lipid binding. GAB1-ΔPH has no effect on EGFR condensate stability (Fig. 5A; Supp. Fig. 5C, 5D), likely because it is not recruited to GUV surfaces. We conclude that PIPs disfavor GAB1 oligomerization and inhibit its ability to reshape EGFR organization through condensate destabilization and GAB1 assembly nucleation.

To elucidate if PIPs regulate biophysical properties of EGFR condensates or GAB1 assemblies, we performed FRAP across conditions and GUV types. As expected, EGFR has a lower mobile fraction in EGFR/GRB2/N-WASP condensates relative to unclustered EGFR on all GUV surfaces (Supp. Fig. 5E-5H; Supp. Fig. 6A). The EGFR mobile fraction decreases further after addition of GAB1-WT but not GAB1-ΔPH (Fig. 5F-5H; Supp. Fig. 6). This effect is observed across GUV types, demonstrating that the diffusion of EGFR within EGFR condensates and GAB1 assemblies is PIP-insensitive (Fig. 5F-5H). In contrast, the mobile fraction of GAB1 is moderately higher on PIP2 and PIP3 GUVs (Fig. 5C), consistent with a reduction in stable GAB1 oligomers. Together, these experiments suggest that PIP2 and PIP3 preferentially bind the monomeric state of GAB1, leading to a reduction in stable GAB1 oligomers on PIP2 and PIP3 GUVs. Thus, PIP2 and PIP3 reduce GAB1’s ability to destabilize EGFR condensates and nucleate new GAB1 assemblies.

**Figure 6:**
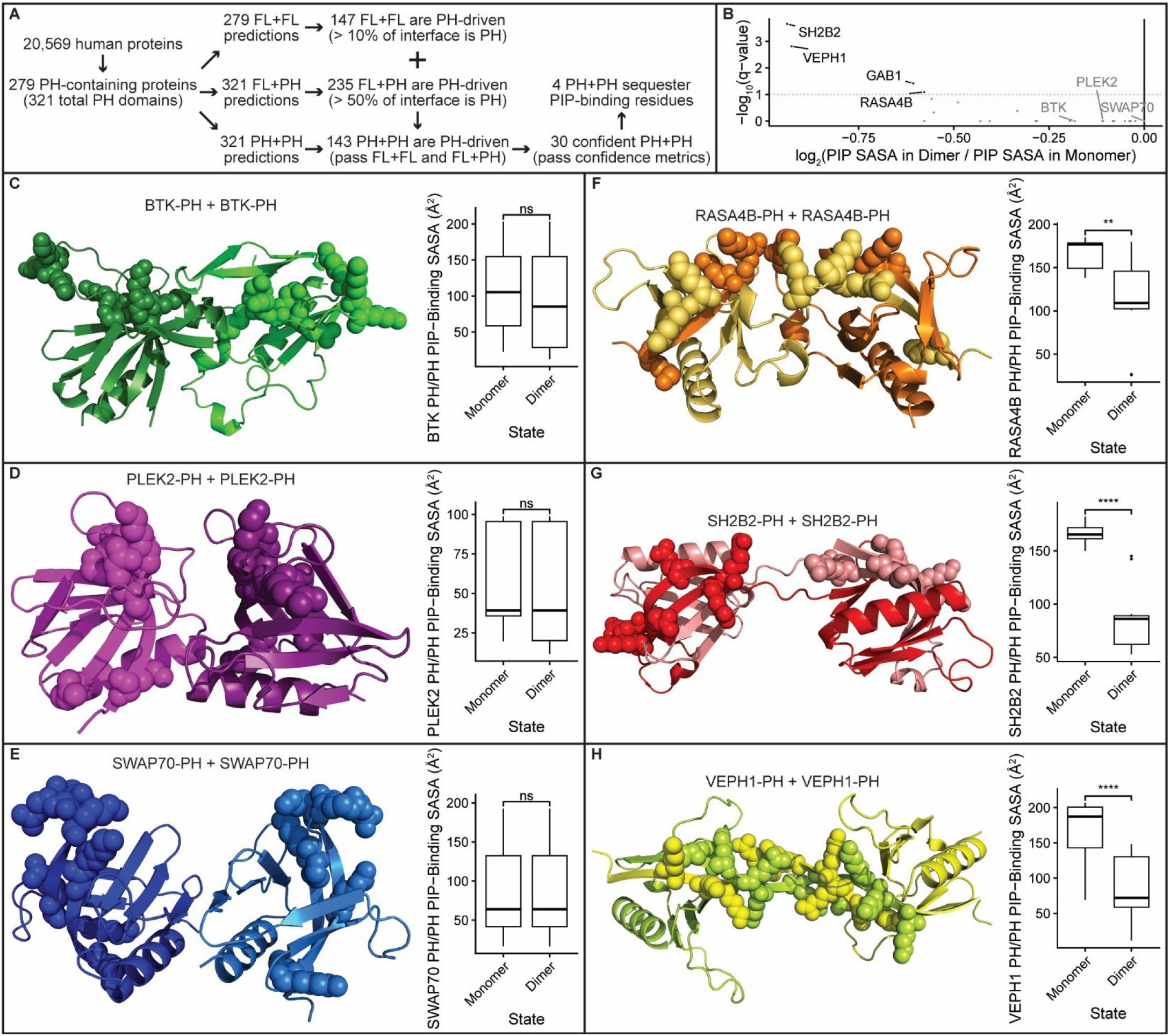
PH domains are predicted to broadly drive dimerization. **(A)** All human PH domains are identified and used for three types of homodimer prediction using AlphaFold: full-length dimers (FL-FL), full-length and PH domain dimers (FL-PH), and PH domain dimers (PH-PH). PH-PH predictions are filtered based on standard scoring metrics and the presence of PH residues in FL-FL and FL-PH interfaces. **(B)** 30 high-confidence PH homodimers are identified and displayed on a volcano plot of their ratio of the solvent-accessible surface area (SASA) of PIP-binding residues in the dimer relative to monomer. This analysis identifies four PH domains, including GAB1, that significantly sequester their PIP-binding residues in the homodimer. **(C-H)** Predicted structures of PH homodimers with the PIP-binding residues displayed as spheres. The box plots display the SASA of these PIP-binding residues in the predicted monomer and dimer structures. BTK (C), PLEK2 (D), and SWAP70 (E) are predicted to form PH homodimers with no significant change in SASA of PIP-binding residues. RASA4B (F), SH2B2 (G), and VEPH1 (H) are predicted to form PH homodimers with significantly reduced SASA of PIP-binding residues. Filtering criteria: FL-FL interface that is PH > 10%, FL-PH interface that is PH > 50%, ipTM > 0.3, pDockQ > 0.23, AlphaJudge Interface Score > 130. Statistics: two-sided Student’s t-test with Benjamini and Hochberg FDR correction (****: p < 0.0001; **: p < 0.01; ns: p > 0.05; n > 5 PIP-binding residues per chain).

### PH domains are predicted to broadly drive dimerization

Our observation that the GAB1 PH domain functions in oligomerization was unexpected. Although PH domains canonically bind PIPs, some PH domains participate in heterotypic and/or homotypic protein interactions^70^. However, of the 321 annotated human PH domains, only the BTK and PDPK1 PH domains have been previously reported to contribute to homodimerization, and both require PIP-containing membranes for this activity^59–62^. To investigate if PH domain-mediated oligomerization generalizes beyond GAB1, we predicted structures of PH domain homodimers *in silico* across the human proteome.

We modeled homodimer structures for all human PH domains in isolation and in their full-length (FL) context with AlphaPulldown^65^, focusing on complexes that are specifically driven by PH-PH interactions (Fig. 6A; Supp. Fig. 7A). This pipeline identified 30 confident PH-PH homodimers (Fig. 6B). Notably, these complexes include the previously described BTK homodimer (Fig. 6C), the pleckstrin protein (PLEK2) where PH domains were first detected (Fig. 6D), and several multifunctional signaling adaptor proteins (Supp. Fig. 7B, 7C). For example, SWAP70 is a scaffolding actin remodeler that is recruited to phagosomes via interactions between its PH domain and PIP2^71^ (Fig. 6E). Unlike GAB1, 26 of the predicted PH domains, including BTK, PLEK2, and SWAP70, are predicted to have their PIP-binding residues exposed in the homodimer state. For these PH domains, lipid binding is expected to promote dimerization, as has been observed experimentally with BTK, which has been shown to not dimerize in solution, even in the presence of PIP headgroups^72^. Neither ITK-PH nor TEC-PH were identified by our approach, in line with their inability to dimerize on PIP3 SLBs^60^. These results indicate that ∼8% of PH domains homodimerize while retaining PIP binding, likely enabling lipid-mediated regulation over oligomerization, downstream signaling, and phase separation.

We also identify four predicted complexes with significantly sequestered PIP-binding residues: GAB1, RASA4B, SH2B2, and VEPH1 (Fig. 6F-6H). Although little is known about PIP binding of these proteins, they all function as adaptors that modulate activity of diverse signaling pathways. RASA4B inhibits RAS/MAPK^73^, SH2B2 interacts with GRB2 downstream of RTKs^74^, and VEPH1 binds EGFR2 to regulate PI3K/AKT^75,76^. These results identify a possible role for PH dimerization in control over signaling networks downstream of RTKs. Notably, no GAB1 paralogs (GAB2, GAB3, GAB4) are predicted to dimerize.

To validate the ability of our *in silico* approach to exclude low-confidence PH homodimers, and to further test effects of PIPs on high-confidence PH homodimers, we performed mass photometry of predicted PH-PH complexes in either PBS alone, or in the presence of IP3 or IP4 (head groups of PIP2 or PIP3, respectively). We selected GAB1-PH-mEGFP as a high-confidence PH homodimer and PLCD1-PH-GST and CYTH3-PH-GST as low-confidence PH homodimers (Supp. Fig. 8A, 8B), which bind PIP2 or PIP3, respectively^77,78^. We also included BTK-PH-SNAP, which is predicted to dimerize *in silico*, but requires PIP-containing membranes to form dimers *in vitro*^60,72,79^. Purification tags were retained to improve protein solubility and increase molecular weight for analysis by mass photometry, which can only detect high-affinity (K_d_ < 200 nM) oligomers that are larger than 30 kDa. GST can dimerize in solution, strengthening the lack of PH-mediated dimerization if no dimers are observed with these constructs, and neither mEGFP nor SNAP form dimers^80–82^.

As expected, GAB1-PH-mEGFP monomers become more abundant in the presence of IP3 and IP4 (Supp. Fig. 8C), confirming that PIP binding stabilizes the monomer. These results provide experimental validation of the AlphaFold-based prediction that lipid binding favors GAB1-PH monomers. These results are also consistent with our observation of fewer GAB1 assemblies on PIP2 or PIP3 GUVs (Fig. 5B). BTK-PH-SNAP, PLCD1-PH-GST, and CYTH3-PH-GST are monomeric in solution (Supp. Fig. 8D-8F).

Despite the high-confidence PH homodimer prediction, previous studies have shown that BTK-PH requires PIP3 SLBs to dimerize^60,72,79^. These results demonstrate successful exclusion of low-confidence PH homodimers (PLCD1 and CYTH3), confirm that GAB1-PH dimers sequester their PIP-binding residues, and demonstrate that the GAB1-PH domain has a higher dimer affinity in solution than the previously characterized BTK-PH domain. Our *in silico* results suggest a general function for PH-mediated dimerization across the human proteome.

## Discussion

In this study, we identify that the GAB1 adaptor protein reshapes EGFR spatial organization through two distinct, independently regulated mechanisms (Fig. 1). GAB1 inhibits GRB2-dependent EGFR phase separation via its long IDR, and it promotes the formation of compositionally distinct assemblies via its PH domain (Supp. Fig. 9). The GAB1 IDR contains many phosphorylation-accessible tyrosines (Supp. Fig. 1F), and we demonstrate that unphosphorylated tyrosines are specifically required to destabilize EGFR condensates. Phosphorylating the IDR prevents EGFR condensate dissolution while having no effect on GAB1 nucleated assemblies (Fig. 2D). Thus, phosphorylation of GAB1 favors EGFR phase separation while dephosphorylation of GAB1 destabilizes large EGFR condensates. These results are strikingly similar to recent work identifying that unphosphorylated tyrosines in PD-1’s ITIM and ITSM inhibit formation of LAT condensates in T-cells^6^. Similar to PD-1, GAB1 contains two ITIMs within its IDR. Unlike PD-1, which inhibits signaling downstream of LAT activation, GAB1 is an adaptor protein that facilitates signaling downstream of EGFR activation. Intriguingly, proteins with at least one ITIM/ITSM in a cytoplasmic IDR are enriched for functions in signaling modulation, immune transduction, and actin remodeling (Fig. 2G), suggesting that this sequence feature might function beyond immune signaling inhibition to more subtly regulate condensate size and signaling in diverse pathways. Synthetic approaches for modulating specific condensates may benefit from incorporating this insight^83^.

The GAB1 PH domain promotes the nucleation of GAB1 assemblies through the formation of stable GAB1 oligomers. Deleting the PH domain has no effect on EGFR condensate dissolution but significantly reduces GAB1 nucleated assemblies (Fig. 2B). Our KMC simulations predict that GAB1 stably oligomerizes in solution (Fig. 3), and we experimentally identify that the PH domain drives GAB1 oligomerization (Fig. 4). These results demonstrate the potential of KMC to discern unknown kinetics and rate constants from phase separation experiments. We further show that PH domain oligomerization can be regulated by membrane lipid composition. We find that lipid binding stabilizes the GAB1 monomer, and that membranes containing PIP2 or PIP3 have fewer GAB1 assemblies (Fig. 5). Thus, increasing PIP2 or PIP3 concentration in the membrane can reduce nucleation of small GAB1 assemblies, but has no direct effect on the larger EGFR condensates.

While we’ve uncovered how distinct molecular features of GAB1 change EGFR spatial organization, we do not fully understand how these changes impact downstream signaling outcomes. EGFR activates several oncogenic signaling networks (e.g. RAS/MAPK, PI3K/AKT, JAK/STAT, PLCγ) via recruitment of distinct adaptor proteins with specific affinities for downstream signaling molecules^8^. Recent work demonstrates that GRB2-dependent EGFR condensates recruit the adaptor protein SOS and enhance RAS signaling^10^. Thus, we predict that destabilizing EGFR condensates would reduce RAS activation, while phosphorylation of GAB1 might increase RAS activation. GAB1 canonically functions to recruit and activate PI3K^35,36^, and we speculate that the GAB1-induced changes in EGFR organization may function to locally switch signaling from RAS to PI3K. However, future experiments measuring RAS and PI3K activity in EGFR condensates and GAB1 assemblies will be required to directly test this hypothesis. The GAB1 PH domain has long been thought to function only as a membrane-targeting domain^13,43,84^. Our unexpected finding that the GAB1 PH domain also drives oligomerization warrants a closer look at prior observations in the literature. Numerous studies have overexpressed GAB1-ΔPH and observed profound effects on downstream signaling. In HeLa cells, the PH domain is required for JNK activation downstream of stimulation with EGF^13^. In Vero cells, the PH domain is required to recruit GAB1 to the membrane and for the phosphorylation of GAB1 tyrosines downstream of stimulation with EGF, but the PH domain is not required for downstream ERK activation^85^. Notably, the GAB1 PIP-binding residues are in the predicted homodimer interface (Fig. 4I). Thus, mutating these residues might reduce recruitment of GAB1 to membranes due to both decreased PIP binding and disrupted oligomerization^43^. While the PH domain is essential for mediating downstream signaling, more careful mutation studies will be required to determine the specific effects of GAB1 PH oligomerization and GAB1 lipid binding in cells. Additionally, while the GAB1 AlphaFold model provides insight into a potential homodimer interface, our experimental data show that the GAB1 PH domain forms higher-order oligomers. Future experiments will be required to identify the intermolecular PH-PH interactions underlying oligomer formation.

We identify that ∼10% of human PH domains are predicted to broadly drive dimerization across diverse human proteins (Fig. 6). Many of these proteins, including all four that significantly sequester their PIP-binding residues, are cytoplasmic adaptors that function downstream of RTKs to regulate distinct signaling pathways. Although homodimers function at every level of signal transduction^86^, only two examples of PH-mediated homodimerization have previously been described, and both require PIP-containing membranes^59–62^. We hypothesize that the three “GAB1-like” PH domains (SH2B2, VEPH1, RASA4B) form dimers in the absence of PIP-containing membranes. However, these proteins are not as well-characterized as GAB1. SH2B2 interacts with GRB2 downstream of RTK activation^74^, and its PH domain is required to inhibit JAK/STAT signaling^87^. VEPH1 is bound by EGFR2 and regulates PI3K/AKT signaling^75,76^, but it is unknown if its PH domain is required for this function. RASA4B is activated by Ca^2+^ and inhibits RAS signaling^73^, but its PH domain has not been characterized. While VEPH1 and RASA4B homodimers have not been described, SH2B2 is reported to dimerize via a distinct dimerization domain and SH2 domains^88,89^. The binding interface in our FL+FL predicted complex contains residues from these previously identified domains, in addition to PH domain residues that have not been previously implicated in dimer formation. Future work will be required to determine if PH domain dimerization or oligomerization is important for how these adaptor proteins control signaling downstream of RTKs and potentially under regulation by PIPs.

Although phosphorylated EGFR has an intrinsic capacity to phase separate with specific adaptor proteins, our findings demonstrate that micron-sized condensates are not necessarily the default organizational state of this receptor. We uncovered multiple mechanisms through which GAB1 actively modulates EGFR phase separation and reshapes receptor organization at membranes. These findings add to a growing body of work uncovering the diverse control parameters cells use to regulate phase separation and restrict condensate size. Posttranslational modifications of scaffold molecules can weaken or strengthen the multivalent interactions driving phase separation^90^. Active processes, including enzymatic activity and transcription, can limit condensate size by changing component concentration or creating gradients^91^. Charge asymmetry between molecules can cause electrostatic effects that limit condensate size and lead to microphase separation^92^. We are only beginning to understand how cells control these parameters to create condensates with specific sizes, emergent properties, and functions. In the context of receptor tyrosine kinase signaling, further mechanistic understanding of how condensate formation, size, emergent properties, and destabilization influence the activities of distinct downstream pathways will better inform development of therapeutics.

## Acknowledgements

We thank members of the Case lab for insightful suggestions; members of the Vos lab for insect cell culture reagents and expertise; Heather Amoroso and Riley Abrashoff of the Koch Institute’s Robert A. Swanson Biotechnology Center; Cassandra Rogers of the W.M. Keck Innovation Center; and Ky Lowenhaupt of the Biophysical Instrumentation Facility. This work was supported by the following grants to L.B.C.: an NSF CAREER Award (#2440518), the Searle Scholars Program (#SSP-2022-113), and the Royal G. and Mae H. Westaway Family Memorial Fund. M.F.U. was partially supported by a postdoctoral fellowship from the Koch Institute’s Ludwig Center. D.E.T. was partially supported by a postdoctoral fellowship from the National Cancer Center. D.D.B. was partially supported by funding from the MIT Undergraduate Research Opportunities Program. A.A.K. acknowledges the kind support of the Koerner family through the Michael and Sonja Endowed Chair. This paper was typeset with the *bioRxiv* word template by @Chrelli: www.github.com/chrelli/bioRxiv-word-template.

## Author contributions

M.F.U. and L.B.C conceived the overall project. M.F.U., D.E.T., and H.C. designed individual experiments. M.F.U. and K.G.S. cloned constructs. M.F.U., D.E.T., and K.G.S. purified proteins. M.F.U. and D.E.T. performed SLB and mass photometry experiments. M.F.U. performed cell experiments. M.F.U. and D.D.B. performed droplet experiments. D.E.T. performed GUV and in silico dimerization experiments. H.C. and A.A.K. developed and simulated the KMC models. M.F.U., D.E.T., H.C., A.A.K, and L.B.C. analyzed and interpreted data. M.F.U., D.E.T., and L.B.C. wrote the manuscript. All authors read, edited, and approved the manuscript.

## Materials and Methods

### DNA cloning

All plasmids were cloned via Gibson Assembly and verified via whole-plasmid sequencing (Plasmidsaurus). EGFR was derived from full-length EGFR in the Human Kinase ORF Kit (Addgene #1000000014) from William Hahn and David Root^93^. Lck (Addgene #186130) and N-WASP (Addgene #186132) were gifts from Michael Rosen^32^. TEV (Addgene #92414) was a gift from David Waugh^94^. ULP1 (Addgene #64697) was a gift from Hideo Iwai^95^. GRB2 (Addgene #46442) was a gift from Bruce Mayer^96^. GAB1-WT was synthesized by Integrated DNA Technologies and used for cloning of all GAB1 constructs. BTK-PH (Addgene #51463) and PLCD1-PH (Addgene #51407) were gifts from Tamas Balla^97^. CYTH3-PH (Addgene #71379) was a gift from Mark Lemmon^98^. HRV3C (Addgene #162795) was a gift from Gottfried Otting^99^. A list of all plasmids used in this study is provided (Supplemental Table 1).

### Bacterial protein expression

For non-Lck proteins, BL21(DE3) (New England Biolabs # C2527H; EGFR, GRB2, N-WASP, Ulp1, GAB1, HRV3C) or Rosetta(DE3) (Fisher Scientific # 71-400-3; TEV, GAB1-PH, BTK-PH, PLCD1-PH, CYTH3-PH) *E. coli* cells were transformed with plasmid DNA via heat shock, plated onto LB plates with appropriate antibiotics, and incubated at 37 °C overnight. An isolated colony was added to 20 mL of LB with appropriate antibiotics and shaken at 175 rpm and 37 °C overnight. Cells were pelleted via centrifugation at 4,000*g* for 15 minutes at 4 °C, resuspended in 20 mL of LB with appropriate antibiotics, diluted in 4-6 L of LB with appropriate antibiotics, and shaken at 200 rpm and 37 °C until OD_600_ > 0.6 (∼2-3 hours). IPTG (Goldbio #I2481C) was added to a final concentration of 1 mM and flasks were shaken at 200 rpm for 16 hours at 18 °C, then at 100 rpm and 4 °C. Cells were pelleted via centrifugation at 4,000*g* for 40 minutes at 4 °C, and non-PH domains were resuspended in 20 mM Tris pH 8 (Fisher Scientific #BP152-500), 300 mM NaCl (Millipore Sigma #S9888), 10 mM imidazole (Thermo Fisher #396745000), 5% glycerol (Fisher Scientific #BP229-4), and 1 mM PMSF (Millipore Sigma #P7626), while PH domains were resuspended in 20 mM HEPES pH 7.5 (Millipore Sigma #H3375), 300 mM NaCl, 20 mM imidazole, 10% glycerol, and 100 μM PMSF. Resuspended pellets were either immediately used for protein purification, or frozen in liquid nitrogen (Linde #UN1977) and stored at -80 °C.

### Insect cell protein expression

For Lck, Sf9 *S. frugiperda* cells (Thermo Fisher #12659017) were transfected with bacmid DNA and transfection reagent (Millipore Sigma #6365779001) and incubated at 27 °C for 2-4 days to generate V0. V0 was then added to Sf9 cells, which were incubated at 27 °C for 2-5 days and centrifuged at 238*g* for 15 minutes at 4 °C to generate V1. V1 was then added to HighFive *T. ni* cells (Thermo Fisher #B85502), which were incubated at 27 °C for 1-2 days at 100 rpm and centrifuged at 238*g* for 30 minutes at 4 °C. Pellets were resuspended in resuspension buffer, and either immediately used for protein purification, or frozen in liquid nitrogen and stored at -80 °C.

### EGFR purification

Protease inhibitor cocktail (1 μg/mL antipain (Millipore Sigma #A6191), 1 μg/mL benzamidine (Millipore Sigma #B6506), 1 μg/mL leupeptin (Goldbio #L-010-5), 1 μg/mL pepstatin (Millipore Sigma #P4265) and 5 mM BME (VWR #97064-878) were added to resuspended cultures on ice. Cells were lysed via cell disruption (Avestin Emulsiflex-C5) and centrifuged at 50,000*g* for 40 minutes at 4 °C. Clarified lysate was applied to a gravity column (Bio-Rad #7372512) with Ni-NTA resin (Fisher Scientific #PI88222), washed with W1 buffer (20 mM Tris pH 8, 150 mM NaCl, 10 mM imidazole, 5 mM BME), W2 buffer (20 mM Tris pH 8, 150 mM NaCl, 20 mM imidazole, 5 mM BME), and eluted with E buffer (20 mM Tris pH 8, 150 mM NaCl, 500 mM imidazole, 5 mM BME). Ni-NTA eluate was diluted 10-fold in QA buffer (20 mM Tris, 1 mM DTT (Fisher Scientific #50-153-2768)), applied to an FPLC column (Cytiva #28406409) with anion-exchange resin (Cytiva #17094705) on an FPLC (Cytiva AKTA pure), and eluted via a linear gradient of QA buffer to QB buffer (20 mM Tris, 1 M NaCl, 1 mM DTT). Q eluate was centrifuged in a 10 kDa concentrator at 4,000*g* and 4 °C (Millipore Sigma #UFC8010) to a final volume of 2 mL, loaded onto an SD75 size exclusion FPLC column (Cytiva # 28989333), and eluted with SEC buffer (150 mM NaCl, 10% glycerol, 25 mM HEPES pH 7.5, 1 mM BME). SEC eluate was centrifuged in a 10 kDa concentrator at 4,000*g* and 4 °C to a final concentration of 25 μM, frozen in liquid nitrogen, stored at -80 °C, and verified by MALDI-MS.

### Lck purification

Protease inhibitor cocktail and 5 mM BME were added to resuspended cultures on ice. Cells were lysed with Dounce homogenizers and centrifuged at 50,000*g* for 40 minutes at 4 °C. Clarified lysate was applied to a gravity column with Ni-NTA resin, washed with W1 buffer, W2 buffer, and eluted with E buffer. Ni-NTA eluate was rocked with 50 nM TEV at 4 °C overnight, diluted 10-fold in QA buffer, applied to an FPLC column with anion-exchange resin on an FPLC, and eluted via a linear gradient of QA buffer to QB buffer. Q eluate was centrifuged in a 10 kDa concentrator at 4,000*g* and 4 °C to a final volume of 2 mL, loaded onto an SD75 size exclusion FPLC column, and eluted with SEC buffer. SEC eluate was centrifuged in a 10 kDa concentrator at 4,000*g* and 4 °C to a final concentration of 25 μM, frozen in liquid nitrogen, stored at -80 °C, and verified by MALDI-MS.

### TEV purification

Protease inhibitor cocktail and 5 mM BME were added to resuspended cultures on ice. Cells were lysed via cell disruption and centrifuged at 50,000*g* for 40 minutes at 4 °C. Clarified lysate was applied to a gravity column with Ni-NTA resin, washed with W1 buffer, W2 buffer, and eluted with E buffer. Ni-NTA eluate was diluted 10-fold in SA buffer (20 mM HEPES pH 7.5, 1 mM DTT), applied to an FPLC column with cation-exchange resin (Cytiva #17094401) on an FPLC, and eluted via a linear gradient of SA buffer to SB buffer (20 mM HEPES pH 7.5, 1 M NaCl, 1 mM DTT). S eluate was centrifuged in a 10 kDa concentrator at 4,000*g* and 4 °C to a final volume of 2 mL, loaded onto an SD75 size exclusion FPLC column, and eluted with SEC buffer. SEC eluate was centrifuged in a 10 kDa concentrator at 4,000*g* and 4 °C to a final concentration of 25 μM, frozen in liquid nitrogen, stored at - 80 °C, and verified by MALDI-MS.

### N-WASP purification

Protease inhibitor cocktail and 5 mM BME were added to resuspended cultures on ice. Cells were lysed via cell disruption and centrifuged at 50,000*g* for 40 minutes at 4 °C. Clarified lysate was applied to a gravity column with Ni-NTA resin, washed with W1 buffer, W2 buffer, and eluted with E buffer. Ni-NTA eluate was rocked with 50 nM TEV at 4 °C overnight, diluted 10-fold in QA buffer, applied to an FPLC column with anion-exchange resin on an FPLC, and eluted via a linear gradient of QA buffer to QB buffer. Q eluate was diluted 10-fold in SA buffer, applied to an FPLC column with cation-exchange resin on an FPLC, and eluted via a linear gradient of SA buffer to SB buffer. S eluate was centrifuged in a 10 kDa concentrator at 4,000*g* and 4 °C to a final volume of 2 mL, loaded onto an SD75 size exclusion FPLC column, and eluted with SEC buffer. SEC eluate was centrifuged in a 10 kDa concentrator at 4,000*g* and 4 °C to a final concentration of 25 μM, frozen in liquid nitrogen, stored at -80 °C, and verified by MALDI-MS.

### GRB2 purification

Protease inhibitor cocktail and 5 mM BME were added to resuspended cultures on ice. Cells were lysed via cell disruption and centrifuged at 50,000*g* for 40 minutes at 4 °C. Clarified lysate was applied to a gravity column with Ni-NTA resin, washed with W1 buffer, W2 buffer, and eluted with E buffer. Ni-NTA eluate was rocked with 50 nM Ulp1 at 4 °C overnight, diluted 10-fold in QA buffer, applied to an FPLC column with anion-exchange resin on an FPLC, and eluted via a linear gradient of QA buffer to QB buffer. Q eluate was centrifuged in a 10 kDa concentrator at 4,000*g* and 4 °C to a final volume of 2 mL, loaded onto an SD75 size exclusion FPLC column, and eluted with SEC buffer. SEC eluate was centrifuged in a 10 kDa concentrator at 4,000*g* and 4 °C to a final concentration of 25 μM, frozen in liquid nitrogen, stored at -80 °C, and verified by MALDI-MS.

### Ulp1 purification

Protease inhibitor cocktail and 5 mM BME were added to resuspended cultures on ice. Cells were lysed via cell disruption and centrifuged at 50,000*g* for 40 minutes at 4 °C. Clarified lysate was applied to a gravity column with Ni-NTA resin, washed with W1 buffer, W2 buffer, and eluted with E buffer. Ni-NTA eluate was rocked with 50 nM TEV at 4 °C overnight, diluted 10-fold in QA buffer, applied to an FPLC column with anion-exchange resin on an FPLC, and eluted via a linear gradient of QA buffer to QB buffer. Q eluate was centrifuged in a 10 kDa concentrator at 4,000*g* and 4 °C to a final volume of 2 mL, loaded onto an SD75 size exclusion FPLC column, and eluted with SEC buffer. SEC eluate was centrifuged in a 10 kDa concentrator at 4,000*g* and 4 °C to a final concentration of 25 μM, frozen in liquid nitrogen, stored at -80 °C.

### SOS-PRR purification

Protease inhibitor cocktail and 5 mM BME were added to resuspended cultures on ice. Cells were lysed via cell disruption and centrifuged at 50,000*g* for 40 minutes at 4 °C. Clarified lysate was applied to a gravity column with GST resin (Cytiva #17075605), washed with GW buffer (50 mM Tris pH 8, 150 mM NaCl, 1 mM DTT), and eluted with GE buffer (10 mM reduced glutathione (Millipore Sigma #G4251), 50 mM Tris pH 8, 150 mM NaCl, 1 mM DTT). GST eluate was rocked with 50 nM TEV at 4 °C overnight, diluted 10-fold in QA buffer, applied to an FPLC column with anion-exchange resin on an FPLC, and eluted via a linear gradient of QA buffer to QB buffer. Q eluate was centrifuged in a 10 kDa concentrator at 4,000*g* and 4 °C to a final volume of 2 mL, loaded onto an SD75 size exclusion FPLC column, and eluted with SEC buffer. SEC eluate was centrifuged in a 10 kDa concentrator at 4,000*g* and 4 °C to a final concentration of 25 μM, frozen in liquid nitrogen, stored at -80 °C, and verified by MALDI-MS.

### GAB1 purification

Identical conditions were used to purify GAB1-WT, GAB1-ΔPH, GAB1-E, and GAB1-F. Protease inhibitor cocktail and 5 mM BME were added to re-suspended cultures on ice. Cells were lysed via cell disruption and centrifuged at 50,000*g* for 40 minutes at 4 °C. Clarified lysate was applied to a gravity column with Ni-NTA resin, washed with W1 buffer, W2 buffer, and eluted with E buffer. Ni-NTA eluate was rocked with 50 nM TEV at 4 °C overnight, diluted 10-fold in QA buffer, applied to an FPLC column with anion-exchange resin on an FPLC, and eluted via a linear gradient of QA buffer to QB buffer. Q eluate was centrifuged in a 30 kDa concentrator (Millipore Sigma #UFC8030) at 4,000*g* and 4 °C to a final volume of 2 mL, loaded onto an SD200 size exclusion FPLC column (Cytiva # 28989335), and eluted with SEC buffer. SEC eluate was centrifuged in a 30 kDa concentrator at 4,000*g* and 4 °C to a final concentration of 25 μM, frozen in liquid nitrogen, stored at -80 °C, and verified by MALDI-MS.

### GAB1-PRR purification

Protease inhibitor cocktail and 5 mM BME were added to resuspended cultures on ice. Cells were lysed via cell disruption and centrifuged at 50,000*g* for 40 minutes at 4 °C. Clarified lysate was applied to a gravity column with GST resin, washed with GW buffer, and eluted with GE buffer. GST eluate was rocked with 50 nM TEV at 4 °C overnight, diluted 10-fold in QA buffer, applied to an FPLC column with anion-exchange resin on an FPLC, and eluted via a linear gradient of QA buffer to QB buffer. Q eluate was centrifuged in a 10 kDa concentrator at 4,000*g* and 4 °C to a final volume of 2 mL, loaded onto an SD75 size exclusion FPLC column, and eluted with SEC buffer. SEC eluate was centrifuged in a 10 kDa concentrator at 4,000*g* and 4 °C to a final concentration of 25 μM, frozen in liquid nitrogen, stored at - 80 °C, and verified by MALDI-MS.

### GAB1-IDR493 purification

Protease inhibitor cocktail and 5 mM BME were added to resuspended cultures on ice. Cells were lysed via cell disruption and centrifuged at 50,000*g* for 40 minutes at 4 °C. Clarified lysate was applied to a gravity column with Ni-NTA resin, washed with W1 buffer, W2 buffer, and eluted with E buffer. Ni-NTA eluate was rocked with 50 nM Ulp1 at 4 °C overnight, diluted 10-fold in QA buffer, applied to an FPLC column with anion-exchange resin on an FPLC, and eluted via a linear gradient of QA buffer to QB buffer. Q eluate was centrifuged in a 10 kDa concentrator at 4,000*g* and 4 °C to a final volume of 2 mL, loaded onto an SD75 size exclusion FPLC column, and eluted with SEC buffer. SEC eluate was centrifuged in a 10 kDa concentrator at 4,000*g* and 4 °C to a final concentration of 25 μM, frozen in liquid nitrogen, stored at -80 °C, and verified by MALDI-MS.

### GAB1-PH purification

Protease inhibitor cocktail, 5 mM BME, and 0.5% Triton X-100 (Millipore Sigma #X100) were added to resuspended cultures on ice. Cells were lysed via cell disruption and centrifuged at 50,000*g* for 40 minutes at 4 °C. Clarified lysate was applied to a gravity column with Ni-NTA resin, washed with W1 buffer, W2 buffer, and eluted with E buffer. Ni-NTA eluate was rocked with 50 nM Ulp1 at 4 °C overnight, centrifuged in a 10 kDa concentrator at 4,000*g* and 4 °C to a final volume of 2 mL, loaded onto an SD75 size exclusion FPLC column, and eluted with SEC buffer. SEC eluate was centrifuged in a 10 kDa concentrator at 4,000*g* and 4 °C to a final concentration of 25 μM, frozen in liquid nitrogen, stored at -80 °C, and verified by MALDI-MS.

### BTK-PH purification

Protease inhibitor cocktail and 5 mM BME were added to resuspended cultures on ice. Cells were lysed via cell disruption and centrifuged at 50,000*g* for 40 minutes at 4 °C. Clarified lysate was applied to a gravity column with Ni-NTA resin, washed with W1 buffer, W2 buffer, and eluted with E buffer. Ni-NTA eluate was rocked with 50 nM HRV3C at 4 °C overnight, diluted 10-fold in SA buffer, applied to an FPLC column with cation-exchange resin on an FPLC, and eluted via a linear gradient of SA buffer to SB buffer. S eluate was centrifuged in a 10 kDa concentrator at 4,000*g* and 4 °C to a final volume of 2 mL, loaded onto an SD75 size exclusion FPLC column, and eluted with SEC buffer. SEC eluate was centrifuged in a 10 kDa concentrator at 4,000*g* and 4 °C to a final concentration of 25 μM, frozen in liquid nitrogen, stored at -80 °C, and verified by MALDI-MS.

### HRV3C purification

Protease inhibitor cocktail and 5 mM BME were added to resuspended cultures on ice. Cells were lysed via cell disruption and centrifuged at 50,000*g* for 40 minutes at 4 °C. Clarified lysate was applied to a gravity column with Ni-NTA resin, washed with W1 buffer, W2 buffer, and eluted with E buffer. Ni-NTA eluate was diluted 10-fold in SA buffer, applied to an FPLC column with cation-exchange resin on an FPLC, and eluted via a linear gradient of SA buffer to SB buffer. S eluate was centrifuged in a 10 kDa concentrator at 4,000*g* and 4 °C to a final volume of 2 mL, loaded onto an SD75 size exclusion FPLC column, and eluted with SEC buffer. SEC eluate was centrifuged in a 10 kDa concentrator at 4,000*g* and 4 °C to a final concentration of 25 μM, frozen in liquid nitrogen, stored at -80 °C, and verified by MALDI-MS.

### PLCD1-PH purification

Protease inhibitor cocktail and 5 mM BME were added to resuspended cultures on ice. Cells were lysed via cell disruption and centrifuged at 50,000*g* for 40 minutes at 4 °C. Clarified lysate was applied to a gravity column with GST resin, washed with GW buffer, and eluted with GE buffer. GST eluate was diluted 10-fold in SA buffer, applied to an FPLC column with cation-exchange resin on an FPLC, and eluted via a linear gradient of SA buffer to SB buffer. S eluate was centrifuged in a 10 kDa concentrator at 4,000*g* and 4 °C to a final volume of 2 mL, loaded onto an SD75 size exclusion FPLC column, and eluted with SEC buffer. SEC eluate was centrifuged in a 10 kDa concentrator at 4,000*g* and 4 °C to a final concentration of 25 μM, frozen in liquid nitrogen, stored at -80 °C, and verified by MALDI-MS.

### CYTH3-PH purification

Protease inhibitor cocktail and 5 mM BME were added to resuspended cultures on ice. Cells were lysed via cell disruption and centrifuged at 50,000*g* for 40 minutes at 4 °C. Clarified lysate was applied to a gravity column with GST resin, washed with GW buffer, and eluted with GE buffer. GST eluate was diluted 10-fold in SA buffer, applied to an FPLC column with cation-exchange resin on an FPLC, and eluted via a linear gradient of SA buffer to SB buffer. S eluate was centrifuged in a 10 kDa concentrator at 4,000*g* and 4 °C to a final volume of 2 mL, loaded onto an SD75 size exclusion FPLC column, and eluted with SEC buffer. SEC eluate was centrifuged in a 10 kDa concentrator at 4,000*g* and 4 °C to a final concentration of 25 μM, frozen in liquid nitrogen, stored at -80 °C, and verified by MALDI-MS.

### Protein phosphorylation

20 μM Lck was activated via incubation with 100 μM ATP (Millipore Sigma #A2383) and 50 μM MgCl_2_ (Thermo Fisher #AM9530G) at room temperature for 45 minutes. Activated Lck was then diluted 100-fold in a solution containing 50 μM EGFR or 50 μM GAB1-WT with 50 mM ATP, 20 mM MgCl_2_, 100 mM HEPES pH 7.5, 150 mM NaCl, and 1 mM BME, then incubated at 30 °C for 18-20 hours. The reaction was stopped via incubation on ice, loaded onto an SD75 Increase size exclusion FPLC column (EGFR; Cytiva #29148721) or SD200 Increase size exclusion FPLC column (GAB1; Cytiva #28990944), and eluted with SEC buffer. SEC eluate was centrifuged in a 10 kDa (EGFR) or 30 kDa (GAB1) concentrator at 4,000*g* and 4 °C to a final concentration of 25 μM, frozen in liquid nitrogen, stored at -80 °C, and verified by MALDI-MS.

### Protein fluorescent labeling

BME was removed from purified protein via a desalting column (Cytiva #29048684), eluting with SEC buffer containing no BME. Desalting eluate was incubated with maleimide Alexa dyes in the 488 (Thermo Fisher #A10254), 568 (Thermo Fisher #A20341), or 647 (Thermo Fisher #A20347) channel at ten-fold molar excess and 4 °C overnight. The reaction was stopped via incubation with 1 mM BME, loaded onto a desalting column, eluted with SEC buffer, centrifuged in a concentrator at 4,000*g* and 4 °C to a final concentration of 25 μM, frozen in liquid nitrogen, and stored at -80 °C.

### Glass cleaning

Glass vials (Fisher Scientific #02-991-468) were incubated with 3 M NaOH (Fisher Scientific #AC450580025) at room temperature for 45 minutes, rinsed four times with water, incubated with 5% Hellmanex (Millipore Sigma #Z805939) at 50 °C for 1 hour, rinsed six times with water, dried at 200 °C for 3 hours, and stored at room temperature in a beaker covered with aluminum foil. Glass well plates (Fisher Scientific #NC0536760) were rotated in 5% Hellmanex at 60 rpm for 4 hours at 55 °C, rinsed 25 times with water, dried with Argon gas (Linde #AR 4.8-T), sealed with an adhesive cover (Thermo Fisher #AB0626), and stored at room temperature.

### Small unilamellar vesicles

Hamilton syringes (VWR #60376) were used to make a 4 μmol lipid mixture of 97.9% POPC (Avanti #850457C), 2% Ni-NTA (Avanti #790404C), and 0.1% PEG (Avanti #880230C) in a clean glass vial. Argon gas was applied to evaporate the lipid mixture and the resulting lipid film was incubated at room temperature under vacuum in a desiccator overnight. The mixture was resuspended in 1.5 mL of PBS (Bio-Rad #1610780), vortexed, subjected to Argon gas, and either immediately used for SUV preparation, or frozen in liquid nitrogen and stored at -80 °C. SUVs were generated by subjecting lipid mixtures to 25-30 freeze/thaw cycles of 30 seconds in liquid nitrogen and 4 minutes at 40 °C, centrifuging at 50,000*g* for 45 minutes at 4 °C, applying Argon gas, and storing at 4 °C for up to 2 weeks.

### Supported lipid bilayer generation

Foil was removed above glass wells, which were then washed with 200 μl of water, incubated with 200 μl of 6 M NaOH at 55 °C for 1 hour, incubated again with 200 μl of 6 M NaOH at 55 °C for 1 hour, washed three times with 200 μl of water, washed three times with 200 μl of SLB buffer (50 mM HEPES pH 7.25, 50 mM KCl (Millipore Sigma #P3911), 1 mM TCEP (Fisher Scientific #58-056-11ML)), and left with 200 μl of SLB buffer. 15 μl of SUVs were added to each well and incubated at 50 °C for 15 minutes. 115 μl of liquid was removed from each well, which were then washed three times with 200 μl of SLB buffer and blocked with 100 μl of 1 mg/mL BSA (Millipore Sigma #A8531) in SLB buffer at room temperature for at least 30 minutes. SLBs were considered stable for the remaining day, and protein addition was staggered for consistent imaging timepoints.

### Supported lipid bilayer phase separation

100 μl of liquid was removed from a well and 100 μl of 300 nM EGFR (15% fluorescently labeled) was added and incubated at room temperature in darkness for 2 hours. Next, 100 μl of liquid was removed from the well, which was then washed three times with 200 μl of SLB buffer with BSA before 200 μg/mL glucose oxidase (Millipore Sigma #G2133), 35 μg/mL catalase (Millipore Sigma #C9322), and 10 mM glucose (Mallinckrodt #4912) was added. SLBs were imaged to verify fluidity, then 100 μl of GRB2 and N-WASP in SLB buffer with BSA was added and imaged, reaching steady state after 30 minutes. 1 μM GAB1 (15% fluorescently labeled) was then added and imaged, reaching steady state after 90 minutes.

### Cell culture

MDA-MB-468 (ATCC #HTB-132) cells were cultured in Dulbecco’s Modified Eagle Medium (Fischer Scientific #11-965-118) with 10% FBS (Thermo Fisher #A5256801) and antibiotics at 37 °C with 5% CO_2_ (Linde #CD 50). Media was changed 2-3 times per week and cells were routinely tested for mycoplasm (Fisher Scientific #NC9719283). For serum starvation experiments, cells were cultured in Dulbecco’s Modified Eagle Medium with antibiotics, but no FBS, at 37 °C with 5% CO_2_ for 18 hours then either fixed immediately, or after addition of 100 nM EGF (Thermo Fisher #AF-100-15-1MG). For immunofluorescence experiments, cells were fixed with 4% PFA (Fisher Scientific #AA433689M) in PBS on ice for 20 minutes, washed three times with cold TBS (Bio-Rad #1706435), permeabilized in 0.2% Triton X-100 in TBS on ice for 15 minutes, washed twice with cold TBST (0.1% Tween 20 (Thermo Fisher # PRH5152)), blocked with blocking buffer (1% BSA in TBST) at room temperature for 1 hour, incubated with primary antibodies (Ms-EGFR (Cell Signaling #2239), Rb-GAB1 (Cell Signaling #93804)) in blocking buffer for 1 hour at room temperature, washed three times with TBST, incubated with secondary antibodies (488-Gt-Ms (Fisher Scientific #PIA32723), 647-Dk-Rb (Fisher Scientific #PIA32795TR)) in blocking buffer for 2 hours at room temperature, washed three times with TBST, and washed once with TBS.

### Droplet assays

Wells were washed three times with water and incubated with droplet buffer (50 mM HEPES pH 7.25, 50 mM KCl, 1 mM TCEP, 1 mg/mL BSA, 5% PEG) for 30 minutes at room temperature. 150 nM EGFR (15% fluorescently labeled), 200 μg/mL glucose oxidase, 35 μg/mL catalase, and 10 mM glucose was added and imaged, followed by GRB2 and N-WASP, then 1 μM GAB1 (15% fluorescently labeled).

### Giant unilamellar vesicle generation

Hamilton syringes were used to make 1.25 μmol lipid mixtures for POPC GUVs (97.9% POPC, 2% Ni-NTA, 0.1% PEG), PIP2 GUVs (95.9% POPC, 2% PIP2 (Avanti #850155P), 2% Ni-NTA, 0.1% PEG), and PIP3 GUVs (95.9% POPC, 2% PIP3 (Avanti #850156P), 2% Ni-NTA, 0.1% PEG) in clean glass vials. 20 μl of lipid mixture was then applied to the conductive side of ITO-slides (Nanion #05 1002) and evaporated with Argon gas. A Vesicle Prep Pro (Nanion) was then used to generate GUVs by using a 16 mm O-ring, 250 μl of 200 mM sorbitol, and applying voltage at 37 °C in four steps: 1) 5 Hz for 10 minutes; 2) ramp from 0 V to 3 V at 5 Hz over 5 minutes; 3) 3V at 5 Hz for 2 hours; 4) ramp from 3 V to 0 V for 3 minutes. GUVs were subjected to Argon gas and stored at 4 °C for less than 2 weeks.

### Giant unilamellar vesicle phase separation

Protein addition was staggered for consistent imaging timepoints. 2 μl of GUVs were incubated with 300 nM EGFR (15% fluorescently labeled), 200 μg/mL glucose oxidase, 35 μg/mL catalase, and 10 mM glucose in 100 μl of GUV buffer (50 mM HEPES pH 7.25, 50 mM KCl, 1 mM TCEP, 1 mg/mL BSA) for 30 minutes at room temperature. GUVs were imaged to verify fluidity, then 100 μl of GRB2 and N-WASP in GUV buffer was added and imaged, reaching steady state after 30 minutes. 1 μM GAB1 (15% fluorescently labeled) was then added and imaged, reaching steady state after 90 minutes.

### Supported lipid bilayer microscopy

SLBs were imaged using LASX acquisition software to drive a Leica TIRF-module mounted on a Leica DMi8 with DIC optics equipped with a plan apo 100x/1.47 NA TIRF oil objective and a Quad Band set filter cube for TIRF applications. Illumination was provided by an integrated laser system equipped with multiple laser lines (405 nm-50mw/488 nm-150mw/561 nm-120mw/638 nm-150mW). Images were acquired using a Hamamatsu Flash 4.0 V3 CMOS camera. FRAP was performed by bleaching in the same laser line as the fluorophore of interest.

### Cell culture microscopy

Cells were imaged using LASX acquisition software to drive a Leica DMi8 with DIC optics equipped with a plan apo 100x/1.47 NA TIRF objective. Illumination was provided by an LED8 light source equipped with two interchangeable filter cubes (one for excitation at 391/32, 479/33, 554/24, 638/31; one for excitation at 473/22, 539/24, 641/78, 810/80). Images were acquired using a Hamamatsu Flash 4.0 V3 CMOS camera.

### Droplet assay microscopy

Droplets were imaged using LASX acquisition software to drive a Leica DMi8 with DIC optics equipped with a 100x/1.47 NA TIRF oil objective. Illumination was provided by an LED8 light source equipped with two interchangeable filter cubes (one for excitation at 391/32, 479/33, 554/24, 638/31; one for excitation at 473/22, 539/24, 641/78, 810/80). Images were acquired using a Hamamatsu Flash 4.0 V3 CMOS camera.

### Giant unilamellar vesicle microscopy

GUVs were imaged using Visi-View acquisition software to drive a Visitech VT-iSIM with an Evident Scientific IX83 inverted microscope stand equipped with a 60x/1.40 NA oil objective. Illumination was provided by an integrated laser system equipped with multiple laser lines (404 nm, 488 nm, 561 nm, 640 nm, 730 nm). Images were acquired using a dual PCO edge 9.4BI camera. FRAP was performed by bleaching in the same laser line as the fluorophore of interest.

### Image analysis

All images were analyzed with custom Fiji macros and R scripts. Raw images are available upon request. All code is provided on GitHub: https://github.com/detillman/case_gab1_2026.

### Analysis of EGFR/GRB2 interactors and ITIM/ITSM proteins

Consensus proteins that interact with both EGFR and GRB2 were identified via the Human Protein Atlas^25^. ITIMs and ITSMs were identified across the human proteome using motifs of [SIVL]-X-Y-X-X-[IVL] and [T-X-Y-X-X-[VI], respectively^100^. Cytoplasmic domains of these proteins were identified with DeepTMHMM v1.0^26^. Each residue of these cytoplasmic domains was scored by AIUPred v2.0 and classified as disordered if their AIUPred score was greater than 0.5^27^. Within these cytoplasmic IDRs, phosphorylation accessibility of tyrosines was scored by Musite v1.0 and tyrosines were classified as phosphorylation-accessible if their Musite score was greater than 0.5^28^. GO enrichment analysis was performed with ClusterProfiler v4.12.6^101^, and similar GO terms were reduced based on semantic similarity via rrvgo v1.16.0^102^. All code is provided on GitHub: https://github.com/detillman/case_gab1_2026.

### In silico simulation

A minimal two-dimensional Kinetic Monte Carlo model of the membrane (100 x 100 periodic lattice), coupled to stochastic exchange with a three-dimensional bulk reservoir, was employed. EGFR/GRB2 were modeled as a membrane-bound complex. N-WASP was modeled as a direct multivalent crosslinker: each EGFR/GRB2 could bind up to four N-WASP molecules and each N-WASP molecule could bridge up to five EGFR/GRB2 sites. GAB1 was modeled as a direct monovalent binder with EGFR/GRB2 that could become multivalent via oligomerization. All kinetic parameters were treated as microscopic channel rates in simulation time units, with event propensities obtained by multiplying each rate by the number of eligible events for each configuration. We include local Monte Carlo relaxation moves as coarse-grained annealing steps, with GAB1-rich puncta represented as finite, membrane-associated droplets using a phenomenological self-limiting droplet free energy:

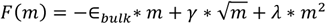

The first term represents the effective drive for puncta growth, the second term penalizes the boundary of a compact two-dimensional membrane-associated droplet, and the final term is an effective coarse-grained penalty that prevents infinite coarsening. We interpret this penalty as a phenomenological representation of finite molecular availability, valence saturation, membrane crowding, and exchange with the soluble pool. This form is motivated by general self-limiting assembly arguments, which imply an excess energetic or kinetic cost that grows superextensively with size^52,53^. All simulation parameters, values, and modifications are provided (Supp. Fig. 3). All code is provided on GitHub: https://github.com/detillman/case_gab1_2026.

### Mass photometry

Proteins were diluted to 200 nM in either PBS, PBS with 1 μM IP3 (Cayman Chemical #10008205), or PBS with 1 μM IP4 (Cayman Chemical #60980). Calibrant solution (Refeyn #MP-CON-41033) was diluted 25-fold in each of these solutions, then diluted 10-fold in PBS and analyzed on a mass photometer (Refeyn OneMP). Proteins were then diluted 10-fold in PBS and analyzed on the mass photometer. DiscoverMP software (Refeyn) was used to identify protein peaks via gaussian fitting and predict molecular weights for these complexes by using results from the calibrant solution in each buffer.

### In silico prediction of pleckstrin homology domain dimers

PH domains and sequences were identified by scraping the Uniprot human proteome UP000005640 for PH domains^103^, as annotated by PROSITE^104^. Homodimers were predicted for full-length proteins (FL-FL), full-length proteins with PH domains (FL-PH), and PH domains (PH-PH) with AlphaPulldown v2.1.8 running on AlphaFold3^63,65^. Interface residues were defined as residues that had C_β_ < 8 Å from a C_β_ on the opposite chain and confidence of predicted complexes was quantified via the Alphafold3 interface-predicted template modelling (ipTM) score, the predicted DockQ score (pDockQ), and the AlphaJudge interface score^63,65,66^. Secondary structure of homodimers was predicted with Define Secondary Structure of Proteins (DSSP) v2.0.4, PIP-binding residues were defined as lysines or arginines on β_1_ or β_2_, or on the loop connecting β_1_ or β_2_^67,68,105^. Predicted structures were rendered in PyMol v3.1.3.1, which was also used to quantify solvent-accessible surface area (SASA) of PIP-binding residues. All code is provided on GitHub: https://github.com/detillman/case_gab1_2026.

---

**Figure S1:**
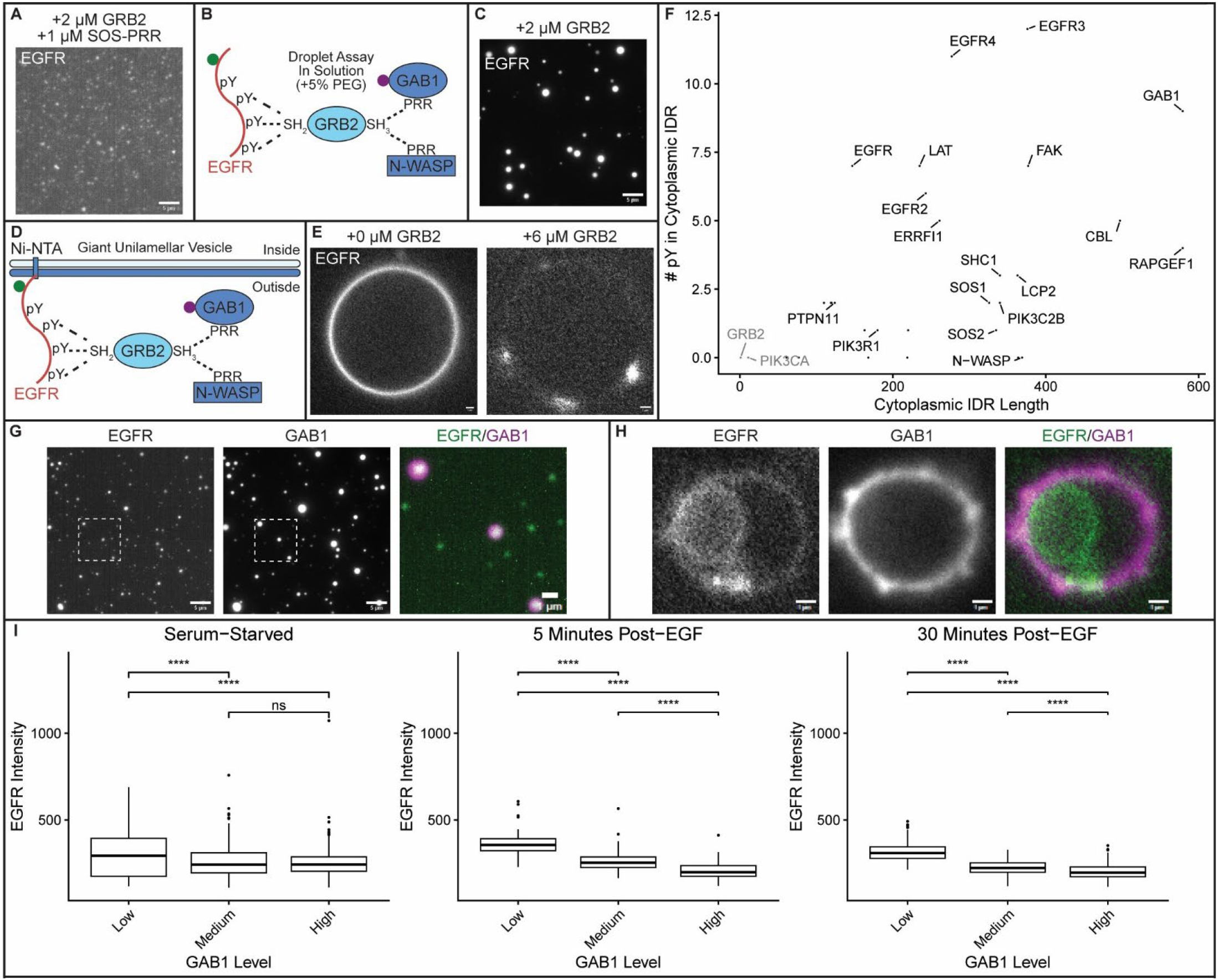
Regulation of EGFR phase separation by GRB2 and GAB1 is conserved across experimental systems. **(A)** EGFR forms condensates with GRB2 and the PRR of SOS on SLBs. **(B)** EGFR is incubated with distinct adaptor proteins in solution. Dashed lines indicate established interactions. **(C)** EGFR phase separates with GRB2 and N-WASP in solution with a crowding agent (5% PEG). **(D)** EGFR is attached to membranes of POPC GUVs with 2% Ni-NTA lipids and incubated with distinct EGFR adaptor proteins. Dashed lines indicate established interactions. **(E)** LEFT: Image of EGFR before adding GRB2 and N-WASP. RIGHT: Image of EGFR after adding GRB2 and N-WASP. **(F)** Scatter plot of the 28 proteins that interact with both EGFR and GRB2, with axes representing cytoplasmic IDR length and number of phosphorylation-accessible tyrosines. **(G)** GAB1 destabilizes EGFR condensates and nucleates distinct GAB1 assemblies in solution with a crowding agent. **(H)** GAB1 destabilizes EGFR condensates and nucleates distinct GAB1 assemblies on GUV membranes. **(I)** Box plots show quantification of EGFR intensity in clusters grouped by low, medium, or high GAB1 intensity during EGF stimulation. Concentrations: 150 nM EGFR, 2 μM GRB2 (A, C, G), 6 μM GRB2 (E, H),1 μM SOS-PRR (A), 1 μM N-WASP (C, E, G, H), 1 μM GAB1 (G, H). SLBs: 97.9% POPC, 2% Ni-NTA, 0.1% DOPE-PEG5000; (C, G): Buffer contains 5% PEG; GUVs: 97.9% POPC, 2% Ni-NTA, 0.1% DOPE-PEG5000. Scale bars: 5 μm; GUV scale bars: 1 μm; Inset scale bars: 1 μm. Statistics: two-sided Student’s t-test (****: p < 0.0001; ns: p > 0.05; n > 86 clusters per condition).

**Figure S2:**
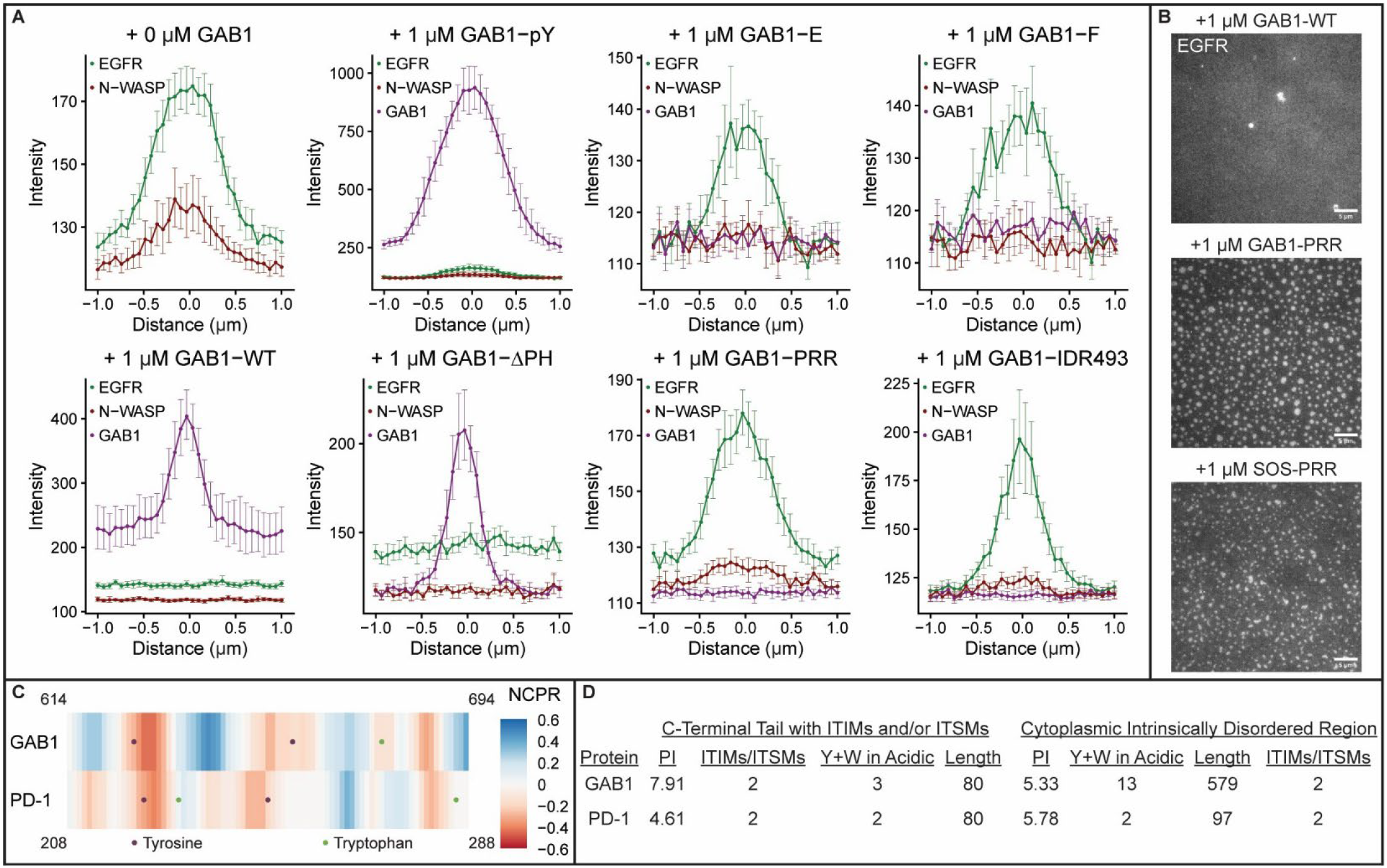
GAB1 nucleates assemblies with compositions distinct from EGFR condensates. **(A)** Plots showing average intensities of different proteins across line scans of puncta on SLBs without GAB1 or after addition of GAB1 variants. **(B)** The PRR of SOS (SOS-PRR) does not destabilize EGFR condensates on SLBs. **(C)** The last 80 residues of GAB1 and PD-1 have similar net charge per residue (NCPR, 5 residue sliding window) patterning, especially around their ITIM/ITSM tyrosines. **(D)** The last 80 residues of GAB1 and PD-1 have similar numbers of tyrosines and tryptophans in acidic patches (NCPR < 0). The cytoplasmic IDRs of GAB1 and PD-1 have similar PIs and per-residue numbers of tyrosines and tryptophans in acidic patches. Line scan error bars: 95% confidence intervals (n > 8 puncta per condition). Concentrations: 150 nM EGFR, 1 μM N-WASP, 6 μM GRB2, 1 μM GAB1 variant, 1 μM SOS-PRR; SLBs: 97.9% POPC, 2% Ni-NTA; 0.1% DOPE-PEG5000. Scale bars: 5 μm.

**Figure S3:**
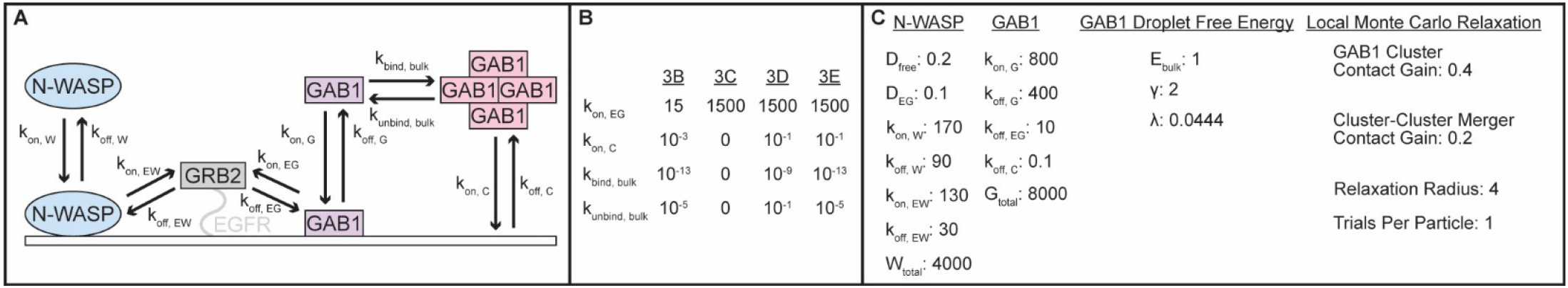
A minimal two-dimensional Kinetic Monte Carlo model with a three-dimensional bulk reservoir reconstitutes EGFR phase separation *in silico*. **(A)** N-WASP solution/membrane exchange (k_on, W_; k_off, W_) and EGFR/GRB2 binding (k_on, EW_; k_off, EW_) are both controlled by two rate constants. Monomeric GAB1 solution/membrane exchange (k_on, G_; k_off, G_) and EGFR/GRB2 binding (k_on, EG_; k_off, EG_) are both controlled by two rate constants. Oligomeric GAB1 solution/membrane exchange (k_on, C_; k_off, C_) and monomer/oligomer turnover (k_bind, bulk_; k_unbind, bulk_) are both controlled by two rate constants. **(B)** Variable parameters across conditions. **(C)** Fixed parameters across conditions. D_free_: diffusion constant for unbound molecules; D_EG_: diffusion constant for EGFR/GRB2 complexes; W_total_: number of N-WASP molecules; G_total_: number of GAB1 molecules; E_bulk_: bulk free energy density per unit of droplet mass; γ: line tension coefficient; λ: superextensive self-limiting coefficient; GAB1 Cluster Contact Gain: contact energy bias for GAB1 monomers in proximity to an existing cluster; Cluster-Cluster Merger Contact Gain: contact energy bias for inter-cluster proximity during trial relaxation; Relaxation Radius: maximum trial displacement sampled during each local relaxation move; Trials Per Particle: Monte Carlo sweep intensity.

**Figure S4:**
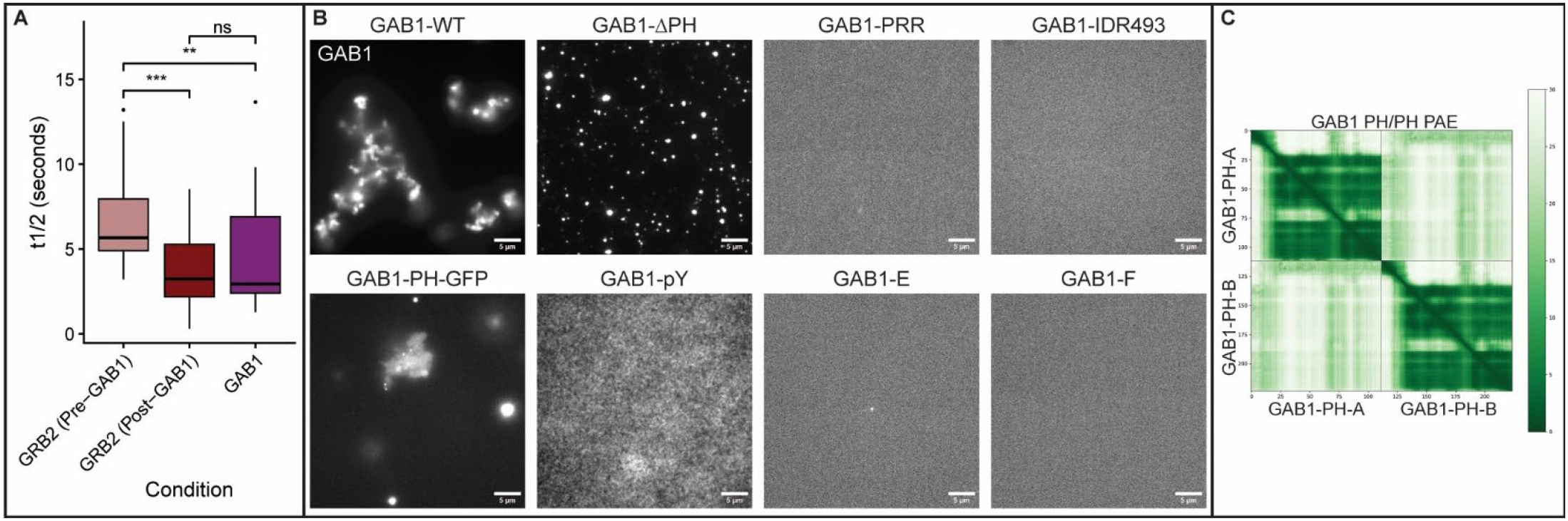
The GAB1 PH domain regulates formation of stable GAB1 oligomers. **(A)** Time to half recovery (t_1/2_) calculated from 4A-4C. **(B)** Fluorescent GAB1 variants added at 1 μM concentration to a solution containing crowding agent (5% PEG). GAB1-WT and GAB1-PH-GFP form oligomeric complexes in solution, while GAB1-ΔPH forms smaller assemblies. No additional GAB1 variants (GAB1-PRR, GAB1-IDR493, GAB1-pY, GAB1-E, GAB1-F) self-associate in this assay. **(C)** Predicted Aligned Error (PAE; Å) identifies contacts between GAB1-PH monomers. Statistics: two-sided Wilcoxon test (****: p < 0.0001; ***: p < 0.001; **: p < 0.01; ns: p > 0.05; n > 16 condensates per condition). Scale bars: 5 μm.

**Figure S5:**
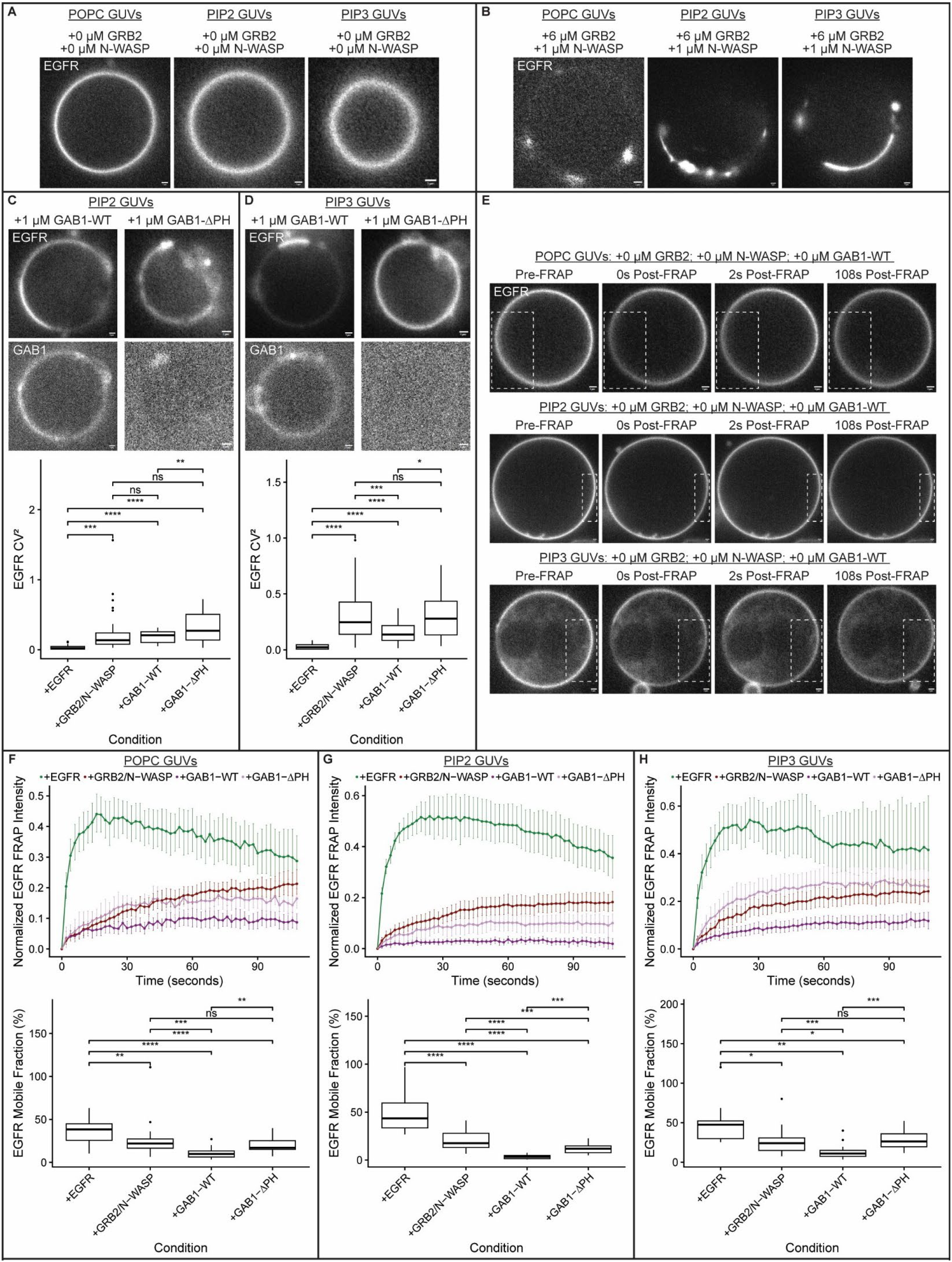
GAB1 destabilizes EGFR condensates on GUVs with phosphatidylinositol phosphate lipids. **(A)** EGFR is added to GUVs containing no phosphatidylinositol phosphate lipids (“POPC GUVs”), 2% phosphatidylinositol 4,5-bisphosphate (“PIP2 GUVs”), or 2% phosphatidylinositol (3,4,5)-trisphosphate (“PIP3 GUVs”). **(B)** Adding GRB2 and N-WASP induces EGFR condensate formation on surfaces of all GUV types. **(C, D)** GAB1-WT, but not GAB1-ΔPH, is recruited to surfaces of PIP2 (C) and PIP3 (D) GUVs, where it nucleates distinct assemblies and destabilizes EGFR condensates, as quantified by the squared coefficient of variation (CV^2^) of EGFR fluorescence along GUV surfaces, which increases with EGFR clustering. **(E)** Representative fluorescence recovery after photobleaching (FRAP) analysis of EGFR on the surface of all GUV types. **(F-H)** FRAP analysis of EGFR across conditions validates formation of EGFR condensates and their destabilization by GAB1-WT (but not GAB1-ΔPH) across POPC (F), PIP2 (G), and PIP3 (H) GUVs. Concentrations: 150 nM EGFR, 1 μM N-WASP, 6 μM GRB2, 1 μM GAB1 variants (C, D); POPC GUVs: 97.9% POPC, 2% Ni-NTA, 0.1% DOPE-PEG5000; PIP2 GUVs: 95.9% POPC, 2% Ni-NTA, 2% PIP2, 0.1% DOPE-PEG5000; PIP3 GUVs: 95.9% POPC, 2% Ni-NTA, 2% PIP3, 0.1% DOPE-PEG5000. Scale bars: 1 μm. Statistics: two-sided Student’s t-test (****: p < 0.0001; ***: p < 0.001; **: p < 0.01; ns: p > 0.05; n > 18 GUVs per condition). FRAP error bars: 95% confidence intervals (n > 10 EGFR condensates per condition).

**Figure S6:**
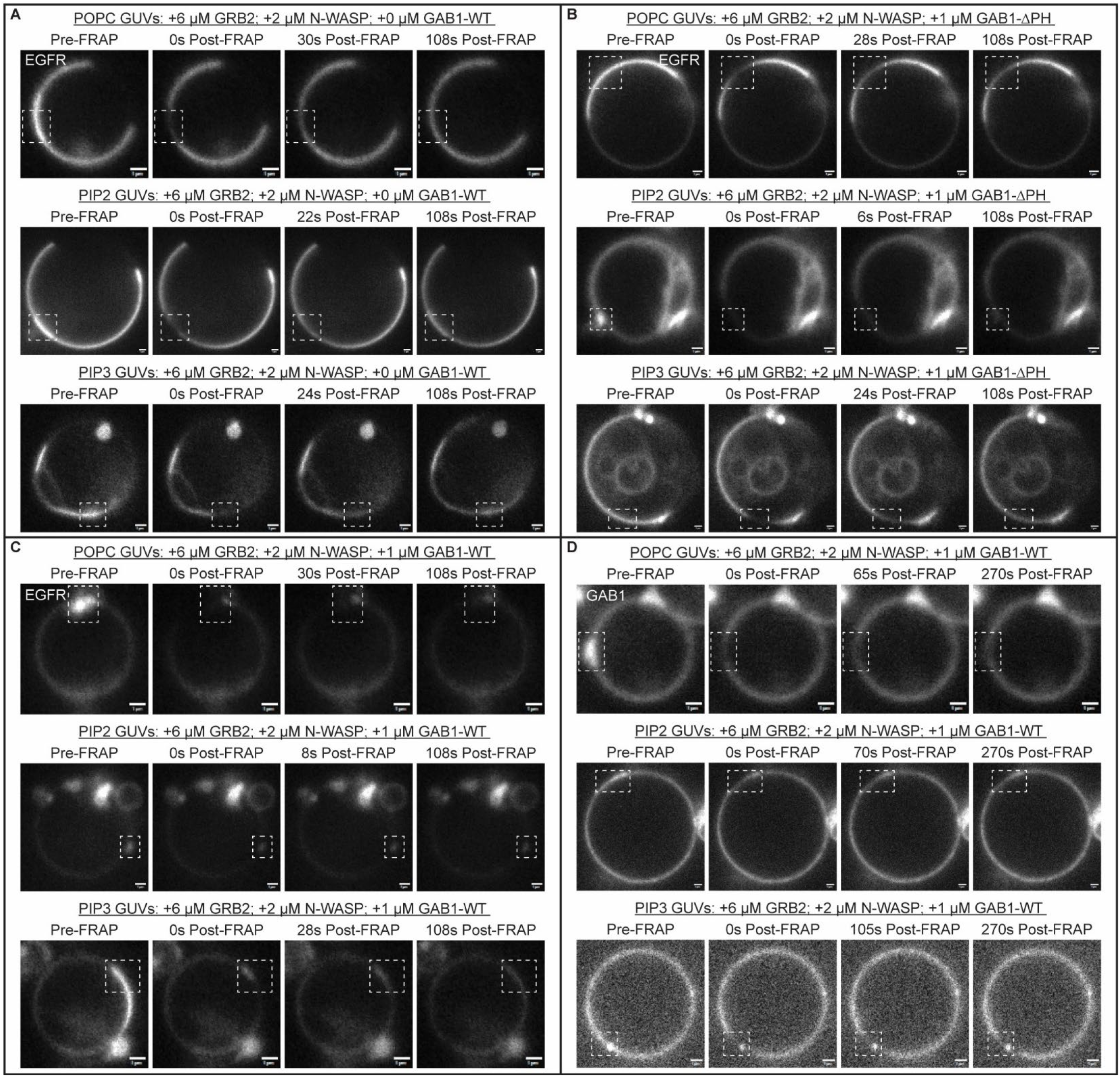
FRAP analysis of EGFR and GAB1 on GUVs. **(A)** Representative images of fluorescence recovery after photobleaching (FRAP) analysis of EGFR after adding GRB2 and N-WASP to EGFR on surfaces of GUVs that have no phosphatidylinositol phosphate lipids (“POPC GUVs”), 2% phosphatidylinositol 4,5-bisphosphate (“PIP2 GUVs”), or 2% phosphatidylinositol (3,4,5)-trisphosphate (“PIP3 GUVs”). **(B)** Representative images of FRAP analysis of EGFR after adding GAB1 without its PH domain (GAB1-ΔPH) to EGFR condensates across GUV types. **(C)** Representative images of FRAP analysis of EGFR after adding full-length GAB1 (GAB1-WT) to EGFR condensates across GUV types. **(D)** Representative images of FRAP analysis of GAB1 after adding full-length GAB1 (GAB1-WT) to EGFR condensates across GUV types. Concentrations: 150 nM EGFR, 1 μM N-WASP, 6 μM GRB2, 1 μM GAB1 variants (B-D); POPC GUVs: 97.9% POPC, 2% Ni-NTA, 0.1% DOPE-PEG5000; PIP2 GUVs: 95.9% POPC, 2% Ni-NTA, 2% PIP2, 0.1% DOPE-PEG5000; PIP3 GUVs: 95.9% POPC, 2% Ni-NTA, 2% PIP3, 0.1% DOPE-PEG5000. Scale bars: 1 μm.

**Figure S7:**
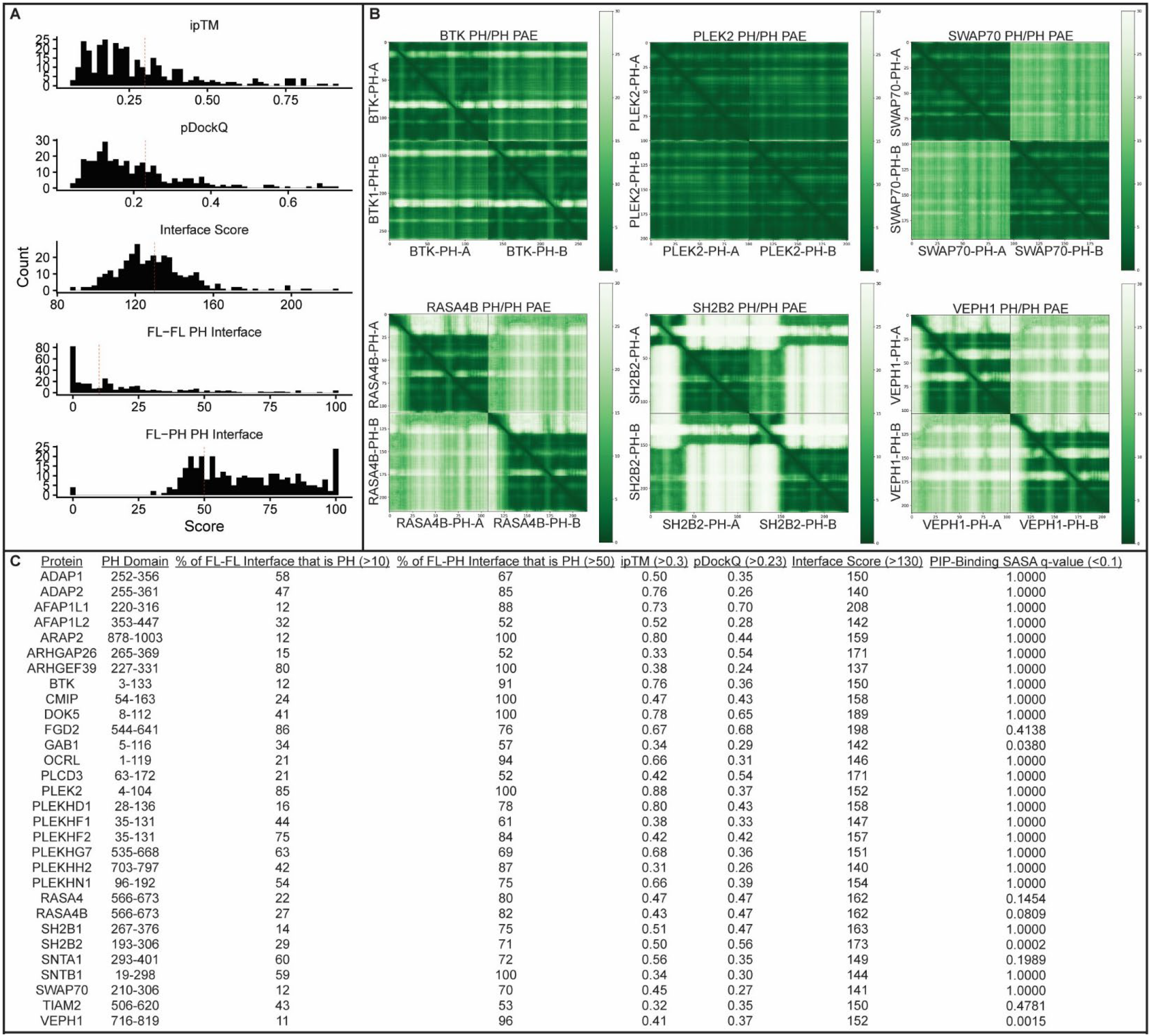
Diverse PH domains are predicted to homodimerize. **(A)** Cutoffs for scoring metrics (dashed red lines) were applied across all predicted PH-PH complexes. **(B)** Predicted Aligned Error (PAE; Å) identifies contacts in PH homodimers. **(C)** Scoring metrics for the thirty high-confidence PH-PH dimers. PH-PH dimers were classified as “high-confidence” if they passed all five scoring metric cutoffs. Filtering criteria: FL-FL interface that is PH > 10%, FL-PH interface that is PH > 50%, ipTM > 0.3, pDockQ > 0.23, AlphaJudge Interface Score > 130.

**Figure S8:**
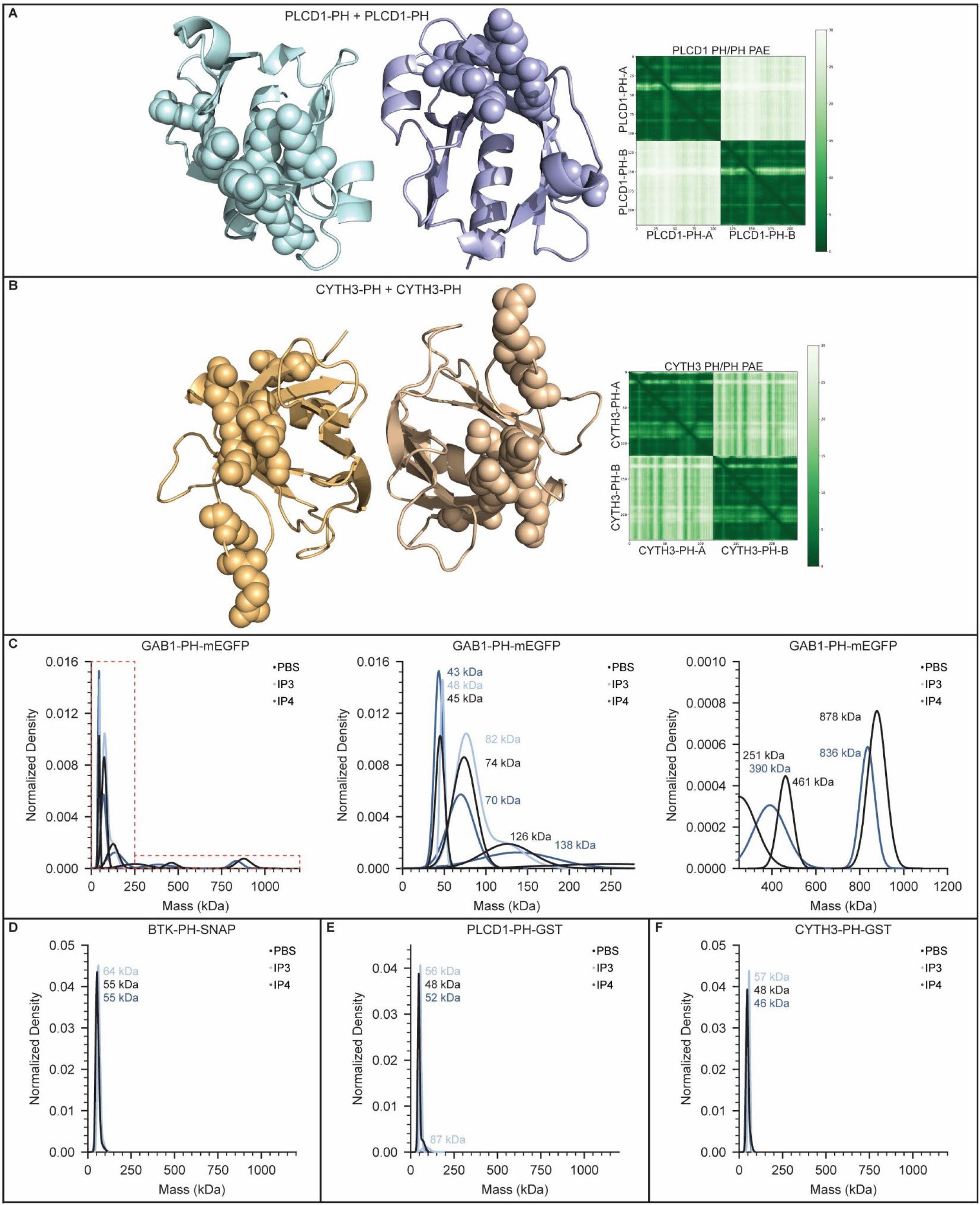
PH domains with low-confidence PH homodimers do not oligomerize in solution. **(A, B)** PLCD1-PH (A) and CYTH3-PH (B) are not predicted to form homodimers. Predicted structures are shown with PIP-binding residues displayed as spheres. Predicted Aligned Error (PAE; Å) identifies few predicted contacts in these homodimers. **(C-F)** Mass photometry measurements to determine the mass of purified PH domains added to solution at 20 nM. IP3 was added to experiments to stabilize the phosphatidylinositol (4,5)-bisphosphate (PIP2) bound conformation. IP4 was added to experiments to stabilize the phosphatidylinositol (3,4,5)-trisphosphate (PIP3) bound conformation. Mass photometry of GAB1-PH-mEGFP (C), BTK-PH-SNAP (D), PLCD1-PH-GST (E), and CYTH3-PH-GST (F). Statistics: two-sided Student’s t-test (ns: p > 0.05; n > 5 PIP-binding residues per chain). Concentrations: 20 nM PH domain construct; Buffer: PBS (“PBS”), PBS with 1 μM IP3 (“IP3”), PBS with 1 μM IP4 (“IP4”).

**Figure S9:**
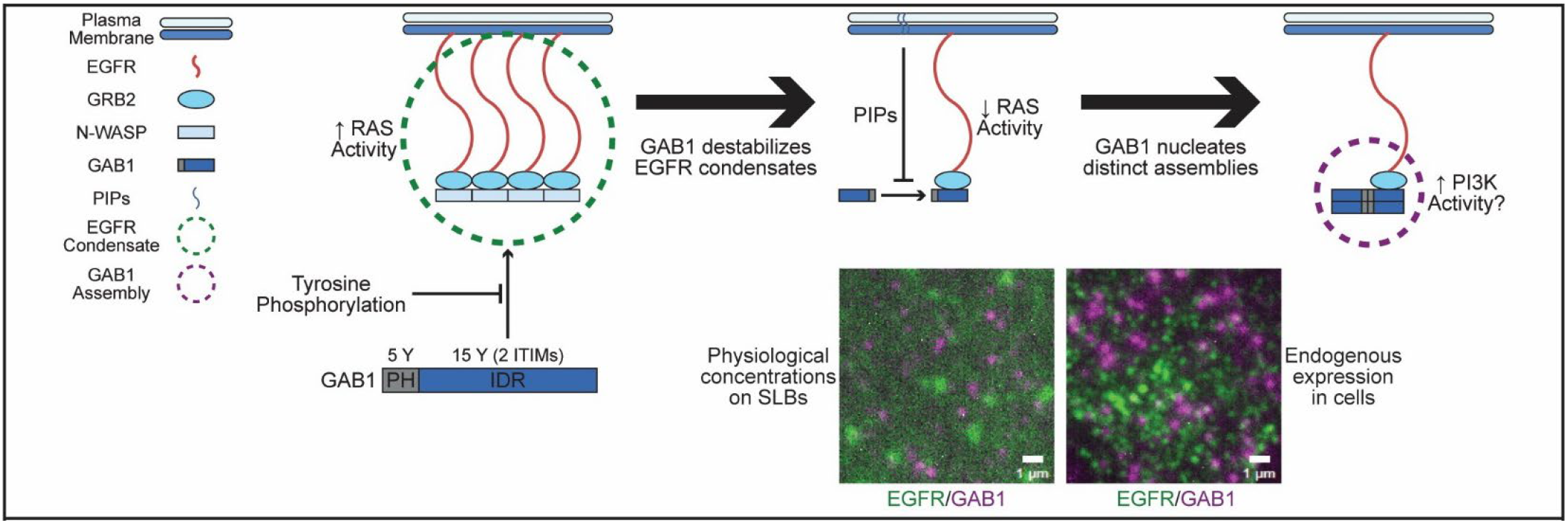
GAxB1 reshapes EGFR organization via PH domain oligomerization and phosphorylation-dependent condensate destabilization. EGFR phase separates with GRB2 and N-WASP on the plasma membrane to increase RAS signaling. GAB1 destabilizes EGFR condensates via unphosphorylated tyrosines (including two ITIMs) in its IDR to decrease RAS activity – phosphorylating GAB1’s tyrosines inhibits this destabilization. The GAB1 PH domain drives oligomerization of GAB1, resulting in nucleation of distinct EGFR assemblies. At physiological protein concentrations, EGFR condensates and GAB1 assemblies coexist on SLBs, leading to an anti-correlation between EGFR and GAB1 intensities (Fig. 1E). This anti-correlation mirrors the organization of EGFR and GAB1 in cells five minutes after EGF stimulation (Fig. 1G).

## References

1. Alberti, S. Phase separation in biology. Current Biology 27, R1097–R1102 (2017).

2. Forman-Kay, J. D., Ditlev, J. A., Nosella, M. L. & Lee, H. O. What are the distinguishing features and size requirements of biomolecular condensates and their implications for RNA-containing condensates? RNA 28, 36–47 (2022).

3. Henninger, J. E. et al. RNA-Mediated Feedback Control of Transcriptional Condensates. Cell 184, 207–225.e24 (2021).

4. Currie, S. L. et al. Quantitative reconstitution of yeast RNA processing bodies. Proceedings of the National Academy of Sciences 120, e2214064120 (2023).

5. Lin, C.-C. et al. Receptor tyrosine kinases regulate signal transduction through a liquid-liquid phase separated state. Molecular Cell 82, 1089–1106.e12 (2022).

6. Sharma, M. et al. Unphosphorylated tyrosines mediate PD-1 inhibition of T cell signaling condensate formation. Science 393, 1145–1151 (2026).

7. Wee, P. & Wang, Z. Epidermal Growth Factor Receptor Cell Proliferation Signaling Pathways. Cancers 9, 52 (2017).

8. Du, Y., Karatekin, F., Wang, W. K., Hong, W. & Boopathy, G. T. K. Cracking the EGFR code: Cancer biology, resistance mechanisms, and future therapeutic frontiers. Pharmacological Reviews 77, 100076 (2025).

9. Needham, S. R. et al. EGFR oligomerization organizes kinase-active dimers into competent signalling platforms. Nat Commun 7, 13307 (2016).

10. Lin, C.-W. et al. A two-component protein condensate of the EGFR cytoplasmic tail and Grb2 regulates Ras activa-tion by SOS at the mem-brane. Proceedings of the National Academy of Sciences 119, e2122531119 (2022).

11. Abulrob, A. et al. Nanoscale imaging of epidermal growth factor receptor clustering: effects of inhibitors. J Biol Chem 285, 3145–3156 (2010).

12. Sachs, M. et al. Essential Role of Gab1 for Signaling by the C-Met Receptor in Vivo. J Cell Biol 150, 1375–1384 (2000).

13. Rodrigues, G. A., Falasca, M., Zhang, Z., Ong, S. H. & Schlessinger, J. A novel positive feedback loop mediated by the docking protein Gab1 and phosphatidylinositol 3-kinase in epidermal growth factor receptor signaling. Mol Cell Biol 20, 1448–1459 (2000).

14. Herbst, R. et al. Daughter of Sevenless Is a Substrate of the Phosphotyrosine Phosphatase Corkscrew and Func-tions during Sevenless Signaling. Cell 85, 899–909 (1996).

15. Hopper, N. A. The Adaptor Protein soc-1/Gab1 Modifies Growth Factor Receptor Output in Caenorhabditis ele-gans. Genetics 173, 163–175 (2006).

16. Itoh, M. et al. Role of Gab1 in Heart, Placenta, and Skin Development and Growth Factor- and Cytokine-Induced Extracellular Signal-Regulated Kinase Mitogen-Activated Protein Kinase Activation. Molecular and Cellular Biology 20, 3695–3704 (2000).

17. Kiyatkin, A. et al. Scaffolding protein Grb2-associated binder 1 sustains epidermal growth factor-induced mito-genic and survival signaling by multiple positive feedback loops. J Biol Chem 281, 19925–19938 (2006).

18. Yamasaki, S. et al. Gab1 is required for EGF receptor signaling and the transformation by activated ErbB2. Onco-gene 22, 1546–1556 (2003).

19. Pérez-Baena, M. J., Cordero-Pérez, F. J., Pérez-Losada, J. & Holgado-Madruga, M. The Role of GAB1 in Cancer. Can-cers 15, 4179 (2023).

20. Case, L. B., Ditlev, J. A. & Rosen, M. K. Regulation of Transmembrane Signaling by Phase Separation. Annu Rev Bio-phys 48, 465–494 (2019).

21. Cheng, X., Ullo, M. F. & Case, L. B. Reconstitution of Phase-Separated Signaling Clusters and Actin Polymerization on Supported Lipid Bi-layers. Frontiers in Cell and Developmental Biology 10, (2022).

22. Bracha, D. et al. Mapping Local and Global Liquid Phase Behavior in Living Cells Using Photo-Oligomerizable Seeds. Cell 175, 1467–1480.e13 (2018).

23. Su, X. et al. Phase separation of signaling molecules promotes T cell receptor signal transduction. Science 352, 595–599 (2016).

24. Banjade, S. & Rosen, M. K. Phase transitions of multivalent proteins can promote clustering of membrane recep-tors. eLife 3, e04123 (2014).

25. Uhlén, M. et al. Tissue-based map of the human proteome. Science 347, 1260419 (2015).

26. Hallgren, J. et al. DeepTMHMM predicts alpha and beta transmembrane proteins using deep neural networks. 2022.04.08.487609 Preprint at 10.1101/2022.04.08.487609 (2022).

27. Erdős, G. & Dosztányi, Z. AIUPred: combining energy estimation with deep learning for the enhanced prediction of protein disorder. Nucleic Acids Res 52, W176–W181 (2024).

28. Wang, D. et al. MusiteDeep: a deep-learning based webserver for protein post-translational modification site prediction and visualization. Nucleic Acids Res 48, W140–W146 (2020).

29. Citri, A. & Yarden, Y. EGF–ERBB signalling: towards the systems level. Nat Rev Mol Cell Biol 7, 505–516 (2006).

30. Sun, S., GrandPre, T., Limmer, D. T. & Groves, J. T. Kinetic frustration by limited bond availability controls the LAT protein condensation phase transition on membranes. Science Advances 8, eabo5295 (2022).

31. Wang, H.-Y. et al. Coupling of protein condensates to ordered lipid domains determines functional membrane organization. Science Advances 9, eadf6205 (2023).

32. Case, L. B., De Pasquale, M., Henry, L. & Rosen, M. K. Synergistic phase separation of two pathways promotes in-tegrin clustering and nascent adhesion formation. eLife 11, e72588 (2022).

33. Kumar, A., Tanaka, K. & Schwartz, M. A. Focal adhesion-derived liquid-liquid phase separations regulate mRNA translation. eLife 13, RP96157 (2025).

34. Lea, N. E. & Case, L. B. Kinase condensates enrich ATP and trigger au-tophosphorylation. Cell Reports 45, (2026).

35. Liu, Y. & Rohrschneider, L. R. The gift of Gab. FEBS Letters 515, 1–7 (2002).

36. Gu, H. & Neel, B. G. The ‘Gab’ in signal transduction. Trends in Cell Biology 13, 122–130 (2003).

37. Tulpule, A. et al. Kinase-mediated RAS signaling via membraneless cytoplasmic protein granules. Cell 184, 2649–2664.e18 (2021).

38. Zhu, T. et al. Phase separation underlies signaling activation of oncogenic NTRK fusions. Proceedings of the National Academy of Sciences 120, e2219589120 (2023).

39. Wiśniewski, J. R., Hein, M. Y., Cox, J. & Mann, M. A “Proteomic Ruler” for Protein Copy Number and Concentration Estimation without Spike-in Standards *. Molecular & Cellular Proteomics 13, 3497–3506 (2014).

40. Filmus, J., Pollak, M. N., Cailleau, R. & Buick, R. N. MDA-468, a human breast cancer cell line with a high number of epidermal growth factor (EGF) receptors, has an amplified EGF receptor gene and is growth inhibited by EGF. Biochem-ical and Biophysical Research Communications 128, 898–905 (1985).

41. Yarden, Y. & Sliwkowski, M. X. Untangling the ErbB signalling network. Nat Rev Mol Cell Biol 2, 127–137 (2001).

42. Mudumbi, K. C. et al. Distinct interactions stabilize EGFR dimers and higher-order oligomers in cell membranes. Cell Reports 43, 113603 (2024).

43. Maroun, C. R. et al. A Conserved Inositol Phospholipid Binding Site within the Pleckstrin Homology Domain of the Gab1 Docking Protein Is Required for Epithelial Morphogenesis*. Journal of Biological Chemistry 274, 31719–31726 (1999).

44. Singh, N. et al. Redefining the specificity of phosphoinositide-binding by human PH domain-containing proteins. Nature Communications 12, 4339 (2021).

45. Holehouse, A. S. & Kragelund, B. B. The molecular basis for cellular function of intrinsically disordered protein regions. Nat Rev Mol Cell Biol 25, 187–211 (2024).

46. McDonald, C. B. et al. Bivalent binding drives the formation of the Grb2-Gab1 signaling complex in a noncoopera-tive manner. FEBS J 279, 2156–2173 (2012).

47. Houtman, J. C. D. et al. Oligomerization of signaling complexes by the multipoint binding of GRB2 to both LAT and SOS1. Nat Struct Mol Biol 13, 798–805 (2006).

48. Gallivan, J. P. & Dougherty, D. A. Cation-π interactions in structural biology. Proceedings of the National Academy of Sciences 96, 9459–9464 (1999).

49. Lin, Y., Currie, S. L. & Rosen, M. K. Intrinsically disordered sequences enable modulation of protein phase separa-tion through distributed tyrosine motifs. J Biol Chem 292, 19110–19120 (2017).

50. Holgado-Madruga, M., Emlet, D. R., Moscatello, D. K., Godwin, A. K. & Wong, A. J. A Grb2-associated docking protein in EGF- and insulin-receptor signalling. Nature 379, 560–564 (1996).

51. Agazie, Y. M. & Hayman, M. J. Development of an Efficient “Substrate-trapping” Mutant of Src Homology Phos-photyrosine Phosphatase 2 and Identification of the Epidermal Growth Factor Receptor, Gab1, and Three Other Pro-teins as Target Substrates*. Journal of Biological Chemistry 278, 13952–13958 (2003).

52. Bollinger, J. A. & Truskett, T. M. Fluids with competing interactions. II. Validating a free energy model for equilib-rium cluster size. The Journal of Chemical Physics 145, 064903 (2016).

53. Hagan, M. F. & Grason, G. M. Equilibrium mechanisms of self-limiting assembly. Rev. Mod. Phys. 93, 025008 (2021).

54. Bartolucci, G., Haugerud, I. S., Michaels, T. C. & Weber, C. A. The inter-play between biomolecular assembly and phase separation. eLife 13, RP93003 (2026).

55. Haugerud, I. S., Bartolucci, G., Braun, D. & Weber, C. A. Theory for sequence selection via phase separation and oligomerization. Proceedings of the National Academy of Sciences 123, e2422829123 (2026).

56. Beutel, O., Maraspini, R., Pombo-García, K., Martin-Lemaitre, C. & Honigmann, A. Phase Separation of Zonula Oc-cludens Proteins Drives Formation of Tight Junctions. Cell 179, 923–936.e11 (2019).

57. Kar, M. et al. Phase-separating RNA-binding proteins form heterogeneous distributions of clusters in subsaturat-ed solutions. Proceedings of the National Academy of Sciences 119, e2202222119 (2022).

58. Wu, D. & Piszczek, G. Measuring the affinity of protein-protein inter-actions on a single-molecule level by mass photometry. Analytical Biochemistry 592, 113575 (2020).

59. Ziemba, B. P., Pilling, C., Calleja, V., Larijani, B. & Falke, J. J. The PH Domain of Phosphoinositide-Dependent Ki-nase-1 Exhibits a Novel, Phospho-Regulated Monomer–Dimer Equilibrium with Important Implications for Kinase Domain Activation: Single-Molecule and Ensemble Studies. Biochemistry 52, 4820–4829 (2013).

60. Chung, J. K. et al. Switch-like activation of Bruton’s tyrosine kinase by membrane-mediated dimerization. Proceed-ings of the National Academy of Sciences 116, 10798–10803 (2019).

61. Chowdhury, S. et al. E41K mutation activates Bruton’s tyrosine kinase by stabilizing an inositol hexakisphos-phate-dependent invisible dimer. Journal of Biological Chemistry 300, (2024).

62. Eisen, T. J. et al. Conditional requirement for dimerization of the mem-brane-binding module for BTK signaling in lymphocyte cell lines. Science Signaling 18, eado1252 (2025).

63. Abramson, J. et al. Accurate structure prediction of biomolecular interactions with AlphaFold 3. Nature 630, 493–500 (2024).

64. Chakravarty, D., Lee, M. & Porter, L. L. Proteins with alternative folds reveal blind spots in AlphaFold-based pro-tein structure prediction. Current Opinion in Structural Biology 90, 102973 (2025).

65. Molodenskiy, D. et al. AlphaPulldown2—a general pipeline for high-throughput structural modeling. Bioinformat-ics 41, btaf115 (2025).

66. Bryant, P., Pozzati, G. & Elofsson, A. Improved prediction of protein-protein interactions using AlphaFold2. Nat Commun 13, 1265 (2022).

67. Lemmon, M. A. Membrane recognition by phospholipid-binding domains. Nat Rev Mol Cell Biol 9, 99–111 (2008).

68. Le Huray, K. I. P., Wang, H., Sobott, F. & Kalli, A. C. Systematic simulation of the interactions of pleckstrin homology domains with mem-branes. Science Advances 8, eabn6992 (2022).

69. Sun, S., Liu, C., Rodriguez Melendez, D., Yang, T. & Cremer, P. S. Immobilization of Phosphatidylinositides Revealed by Bilayer Leaflet Decoupling. Journal American Chemical Society 142, 13003–13010 (2020).

70. Scheffzek, K. & Welti, S. Pleckstrin homology (PH) like domains - versatile modules in protein-protein interaction platforms. FEBS Lett 586, 2662–2673 (2012).

71. Baranov, M. V. et al. SWAP70 Organizes the Actin Cytoskeleton and Is Essential for Phagocytosis. Cell Reports 17, 1518–1531 (2016).

72. Baraldi, E. et al. Structure of the PH domain from Bruton’s tyrosine kinase in complex with inositol 1,3,4,5-tetrakisphosphate. Structure 7, 449–460 (1999).

73. Lockyer, P. J., Kupzig, S. & Cullen, P. J. CAPRI regulates Ca2+-dependent inactivation of the Ras-MAPK pathway. Current Biology 11, 981–986 (2001).

74. Qian, X., Riccio, A., Zhang, Y. & Ginty, D. D. Identification and Characterization of Novel Substrates of Trk Receptors in Developing Neurons. Neuron 21, 1017–1029 (1998).

75. Shathasivam, P. et al. Human ortholog of Drosophila Melted impedes SMAD2 release from TGF-β receptor I to inhibit TGF-β signaling. Proceedings of the National Academy of Sciences 112, E3000–E3009 (2015).

76. Kollara, A., Burt, B. D., Ringuette, M. J. & Brown, T. J. The adaptor protein VEPH1 interacts with the kinase domain of ERBB2 and impacts EGF signaling in ovarian cancer cells. Cellular Signalling 106, 110634 (2023).

77. Lomasney, J. W. et al. Phosphatidylinositol 4,5-Bisphosphate Binding to the Pleckstrin Homology Domain of Phospholipase C-δ1 Enhances Enzyme Activity*. Journal of Biological Chemistry 271, 25316–25326 (1996).

78. Corbin, J. A., Dirkx, R. A. & Falke, J. J. GRP1 Pleckstrin Homology Domain: Activation Parameters and Novel Search Mechanism for Rare Target Lipid. Biochemistry 43, 16161–16173 (2004).

79. Wang, Q., Pechersky, Y., Sagawa, S., Pan, A. C. & Shaw, D. E. Structural mechanism for Bruton’s tyrosine kinase acti-vation at the cell membrane. Proceedings of the National Academy of Sciences 116, 9390–9399 (2019).

80. Fabrini, R. et al. Monomer−Dimer Equilibrium in Glutathione Transferases: A Critical Re-Examination. Biochemis-try 48, 10473–10482 (2009).

81. Zacharias, D. A., Violin, J. D., Newton, A. C. & Tsien, R. Y. Partitioning of Lipid-Modified Monomeric GFPs into Mem-brane Microdomains of Live Cells. Science 296, 913–916 (2002).

82. Dreyer, R., Pfukwa, R., Barth, S., Hunter, R. & Klumperman, B. The Evolution of SNAP-Tag Labels. Biomacromole-cules 24, 517–530 (2023).

83. Savinov, A. et al. AI-discovered protein fragments as generalizable regulators of biomolecular condensates. 2026.05.08.723928 Preprint at 10.64898/2026.05.08.723928 (2026).

84. Maroun, C. R. et al. The Gab1 PH domain is required for localization of Gab1 at sites of cell-cell contact and epithe-lial morphogenesis down-stream from the met receptor tyrosine kinase. Mol Cell Biol 19, 1784–1799 (1999).

85. Yart, A. et al. A Critical Role for Phosphoinositide 3-Kinase Upstream of Gab1 and SHP2 in the Activation of Ras and Mitogen-activated Protein Kinases by Epidermal Growth Factor *. Journal of Biological Chemistry 276, 8856–8864 (2001).

86. Klemm, J. D., Schreiber, S. L. & Crabtree, G. R. Dimerization as a Regulatory Mechanism in Signal Transduction. Annual Review of Immunology 16, 569–592 (1998).

87. Wakioka, T. et al. APS, an adaptor protein containing Pleckstrin homology (PH) and Src homology-2 (SH2) do-mains inhibits the JAK-STAT pathway in collaboration with c-Cbl. Leukemia 13, 760–767 (1999).

88. Hu, J., Liu, J., Ghirlando, R., Saltiel, A. R. & Hubbard, S. R. Structural Basis for Recruitment of the Adaptor Protein APS to the Activated Insulin Receptor. Molecular Cell 12, 1379–1389 (2003).

89. Dhe-Paganon, S. et al. A phenylalanine zipper mediates APS dimerization. Nat Struct Mol Biol 11, 968–974 (2004).

90. Kim, T. H. et al. Phospho-dependent phase separation of FMRP and CAPRIN1 recapitulates regulation of transla-tion and deadenylation. Science 365, 825–829 (2019).

91. Sorkin, B. & Wingreen, N. S. Accelerated Ostwald Ripening by Chemical Activity. Journal American Chemical Society 147, 44673–44679 (2025).

92. Luo, C. et al. Theory of Condensate Size Control by Molecular Charge Asymmetry. ACS Macro Lett. 14, 1484–1491 (2025).

93. Johannessen, C. M. et al. COT drives resistance to RAF inhibition through MAP kinase pathway reactivation. Nature 468, 968–972 (2010).

94. Raran-Kurussi, S., Cherry, S., Zhang, D. & Waugh, D. S. Removal of Affinity Tags with TEV Protease. in Heterologous Gene Expression in E.coli: Methods and Protocols (ed. Burgess-Brown, N. A.) 221–230 (Springer, New York, NY, 2017). doi:10.1007/978-1-4939-6887-9_14.

95. Guerrero, F., Ciragan, A. & Iwaï, H. Tandem SUMO fusion vectors for improving soluble protein expression and purification. Protein Expression and Purification 116, 42–49 (2015).

96. Machida, K. et al. High-Throughput Phosphotyrosine Profiling Using SH2 Domains. Molecular Cell 26, 899–915 (2007).

97. Várnai, P. & Balla, T. Visualization of Phosphoinositides That Bind Pleckstrin Homology Domains: Calcium- and Agonist-induced Dynamic Changes and Relationship to Myo-[3H]inositol-labeled Phosphoinositide Pools. J Cell Biol 143, 501–510 (1998).

98. Kavran, J. M. et al. Specificity and promiscuity in phosphoinositide binding by pleckstrin homology domains. J Biol Chem 273, 30497–30508 (1998).

99. Abdelkader, E. H. & Otting, G. NT*-HRV3CP: An optimized construct of human rhinovirus 14 3C protease for high-yield expression and fast affinity-tag cleavage. Journal of Biotechnology 325, 145–151 (2021).

100. Liu, H. et al. A Comprehensive Immunoreceptor Phosphotyrosine-based Signaling Network Revealed by Recipro-cal Protein–Peptide Array Screening*. Molecular & Cellular Proteomics 14, 1846–1858 (2015).

101. Xu, S. et al. Using clusterProfiler to characterize multiomics data. Nature Protocols 19, 3292–3320 (2024).

102. Sayols, S. rrvgo: a Bioconductor package for interpreting lists of Gene Ontology terms. microPublication. Biology 2023, (2023).

